# Cross-sectional study of antimicrobial resistance and ecology in gastrointestinal and oral microbial communities of urban Pakistani adults

**DOI:** 10.1101/2022.11.05.515288

**Authors:** Maria Batool, Ciara Keating, Sundus Javed, Arshan Nasir, Muhammad Muddassar, Umer Zeeshan Ijaz

**Author notes:** **Correspondence: Umer Zeeshan Ijaz**, RM3111, Level 3, Advanced Research Centre (ARC), 11 Chapel Lane, Glasgow, G11 6EW, UK. **Disclosure: (Conflict of interest)** Before data analysis and manuscript drafting, AN became an employee of and shareholder in Moderna Inc. **Writing Assistance:** The authors did not use writing assistance services as native English speakers have contributed to the writing of the manuscript. **Author Contributions:** Maria Batool, (Data curation: Equal; Investigation: Equal; Methodology: Equal; Software: Equal; Writing – original draft: Equal; Writing – review & editing: Equal). Ciara Keating, PhD (Formal analysis: Equal; Investigation: Equal; Supervision: Supporting; Writing – original draft: Equal; Writing – review & editing: Equal). Sundus Javed, PhD (Formal Analysis: Supporting; Investigation: Supporting; Writing – review & editing: Supporting; Supervision: Supporting). Arshan Nasir, PhD (Data curation: Equal; Investigation: Equal; Methodology: Equal; Project administration: Supporting; Writing – review & editing: Supporting; Resources: Supporting). Muhammad Muddassar, PhD (Formal analysis: Equal; Project administration: Lead; Writing – review & editing: Supporting; Supervision: Lead). Umer Zeeshan Ijaz, PhD (Conceptualization: Lead; Resources: Lead; Software: Lead; Investigation: Equal; Supervision: Lead; Visualization: Lead; Formal analysis: Equal; Writing – original draft: Equal; Writing – review & editing: Equal; Funding Acquisition: Lead). **Data Transparency Statement:** Sequencing data and anonymised meta-data are available upon request. Some of the scripts are available as part of statistical packages available at http://userweb.eng.gla.ac.uk/umer.ijaz#bioinformatics.

## Abstract

**Background and Aims:** Antimicrobial resistance (AMR) is one of the most serious global public health threats affecting lower-middle-income countries (LMICs) due to lack of awareness, inadequate healthcare and sanitation infrastructure, plus other environmental factors. In this study, we aimed to link microbial assembly and covariates (body mass index, smoking, use of antibiotics) to gut microbiome structure and correlate AMR gene prevalence.

**Methods:** We examined the gastrointestinal and oral microbial profiles of healthy adults in Pakistan through 16S rRNA gene sequencing with a focus on different ethnicities, antibiotic usage, drinking water type, smoking, and other demographic measures. We then utilised a suite of innovative statistical tools, driven by numerical ecology and machine learning, to address the above aims.

**Results:** We observed tap water as the main contributor for development of AMR in the Pakistani cohort. In addition, microbial niche breadth analysis based on null modelling procedures highlighted an aberrant gut microbial signature of smokers with increased age.

**Conclusions:** Drinking water plays a more important role in AMR spread in Pakistan rather than other factors considered. Moreover, covariates such as smoking, and age impact the human microbial community structure in this Pakistani cohort. To the best of our knowledge, this is one of the first studies that provide a snapshot of the microbiomes of healthy individuals in Pakistan and considers AMR profiles with an emphasis on potential sources of AMR prevalence.

**Background and Context:** Pakistan is categorized as a low-and-middle-income country by the World Bank where misuse of antibiotics is widespread, and multidrug resistance is prevalent. Thus, it is imperative that we understand antimicrobial resistance and the drivers of human microbiomes in Pakistan.

**New Findings:** In a healthy Pakistani cohort, individuals that consumed *Tap Water* had almost 6-fold more associations with AMRs. Therefore, drinking water source could be a strong driver in the spread of AMR.

**Limitations:** A limitation is the use of predictive functional profiles. However, shotgun metagenomics may be prohibitively costly for LMICs given the urgent need for AMR surveillance.

**Clinical Research Relevance:** Our research shows strong associations of key microbial taxa with covariates such as age, BMI, and gender. Additionally, we show correlations between specific outlier taxa that are present both in the gut and oral communities, highlighting potential future feasibility for use of the oral microbiome as a proxy to gut dysbiosis in some cases.

**Basic Research Relevance:** We have applied recent advancements in analytical tools to link both AMR prevalence and human microbiome composition with factors such as age, BMI, gender, ethnicity, smoking status, use of antibiotics, and drinking water source. Additionally, we use null modelling to show that the microbial communities are subject to strong environmental pressure and dispersal limitation.

**Lay Summary:** We analysed gut and oral microbes from healthy individuals in Pakistan and found that the potential for antibiotic resistance was increased in those who drank tap water.

## Introduction

The discovery of antibiotics was one of the greatest scientific achievements of the 20^th^ century. Many deadly infectious diseases such as typhoid fever, pneumonia, syphilis and tuberculosis can now be treated using standard antibiotics, improving patient treatment outcomes.^1^ Bacteria, however, use many mechanisms to evade antibiotics giving rise to antimicrobial resistance (AMR).^2^ Overuse of antibiotics, especially in agriculture, livestock, and public health, has fuelled AMR evolution in bacterial pathogens, particularly in Gram-negative bacteria.^3^ AMR has now emerged as one of the top 10 global public health threats of 21^st^ century. It is estimated that in 2019, approximately 4.95 million deaths were associated with bacterial AMR.^4^ A review by the UK Government proposed that by 2050 AMR could kill 10 million people per year worldwide.^5^ Quantifying the global burden of AMR to formulate suitable public health measures is difficult due to the unavailability of high-quality population-based data.^6^ In 2015, the WHO launched the global antimicrobial resistance and use surveillance system (GLASS) program for AMR surveillance at global level.^7^ The direct and indirect impact of AMR is expected to mostly fall on LMICs, due to high infectious disease burden and associated mortality.^8^ Moreover they lag behind in active surveillance systems to monitor and control antibiotic administration to the general public.^9^ Other key contributing factors include the availability of antibiotics without prescription, lack of knowledge, and uncontrolled use of antibiotics in agriculture and livestock.^10^ Pakistan is ranked 5^th^ in the world population, with > 220 million people living in the country, and has a high prevalence of antibiotic resistant bacteria, posing a significant regional and global threat.^11^ The overconsumption and inappropriate usage of antibiotics have been reported by several studies.^11,12,13^ Cephalosporins, tetracycline, macrolides and quinolones are the most prescribed drug groups in Pakistan.^14^ Cephalosporins (3^rd^ generation broad-spectrum antibiotics) are prescribed to 67% of patients in Pakistani hospitals.^12^ Recent situational analysis of AMR rates amongst GLASS-specified pathogens in Pakistan reported greater than 50% resistance rates for 3^rd^ generation broad-spectrum antibiotics including cephalosporins and fluoroquinolones among *Klebsiella pneumoniae* and *Escherichia coli.*^15^ Moreover, a 2020 study also reported that the clinical isolates of *Acinetobacter baumannii* harbored 100% resistance to almost every antibiotic tested, suggesting the extensive occurrence of multi-drug resistant strains of *A. baumannii* in Pakistan.^16^

The human gut is a reservoir for various microbes harboring antibiotic resistance genes, which under certain conditions can cause infectious disease.^17^ The gut microbiota in general are commensal in nature, aiding with digestion, immunity, nutrient acquisition and protection against pathogens.^18, 19^ Exposure to antibiotics can cause dysbiosis (i.e., alteration in the taxonomic composition and overall microbial diversity) in the resident gut microbiota.^20^ Some bacterial species possess intrinsic mechanisms to counteract antibiotics and can become enriched post exposure. On the other hand, gut dysbiosis promotes horizontal gene transfer (HGT) of resistance-conferring genes and fuels the evolution of drug-resistant pathogens and the spread of antibiotic resistance.^21^ It also causes a reduction in the diversity of known beneficial bacterial phyla, such as *Bacteroidetes* and *Firmicutes*, which dominate the gut microbiota of healthy adults.^22^ This is a potential concern as the healthy gut microbiota provides colonization resistance from invading pathogenic bacteria.^23^

The gut is a highly diverse ecosystem of microorganisms. In this environment, on the one hand, a large proportion of the resident microbes are highly stable and present from early life.^24^ While on the other hand, external factors such as diet, age, and disease development can impact microbial composition.^25, 26^ Moreover, gut microbial communities are additionally driven by seemingly random processes such as mutations, births, deaths, and ecological drift.^27^ We can elucidate the contribution of these different factors towards how communities are assembled within the gut environment. This is done by employing null modelling tools where broadly, the goal is to see if the underlying structure is driven by *deterministic* or *stochastic* (random) processes.^28, 29^ The application of these tools to the human gut microbiome is still relatively new.^30^ Studies have found, perhaps unsurprisingly, that strong deterministic forces can shape the human gut microbiome. However, dispersal limitation is also highlighted as a key process.^31^ Stegen et al. (2018) argued that cross-talk between environmental and human microbiomes is required to understand and ultimately manipulate or modulate microbiomes.^30^ One technique in environmental or ecosystem analysis is niche breadth, which refers to the environment or constraints within which species or populations can survive.^32^ This concept has recently been applied to the human gut.^33^ The authors found that specific specialist taxa occupied a niche within the human gut during post-bariatric surgery. It is also known that the human gut microbiome can be highly individual.^25^ However, it is unclear whether these differences are unique or related to individual habits. The application of niche breadth to human gut environments (dictated by *Ethnicity*, *Smoking Status*, *Antibiotics Usage Status*, for example) could help to unravel these factors and identify which microbes are context specific to these key environments.

In our previous work, we performed a characterisation study of the gut and oral microbial communities of healthy individuals in Pakistan.^34^ In this present study, we aim to provide a cross-sectional analysis and interrogate this data further with respect to differences in body mass index (BMI), smoking, the use of antibiotics and other covariates. Specifically, our aims are to i) quantify the prevalence of AMR genes, ii) determine how confounding factors such as BMI, smoking, source of drinking water, dietary habits and antibiotic use contribute to microbial community structure and link taxa to these specific niches and finally iii) to determine what forces are driving microbial community structure (i.e., host conditions within the immediate environment or random processes). To address these aims we have recovered functional profiles of microbial communities using PiCRUSt2 ^35^ and have used KEGG signatures of antimicrobial resistance genes ^36^ to quantify AMR prevalence. We then used *Generalised Linear Latent Variable Models* (GLLVM) to link abundance of microbes as well as AMR KOs to the sources of variation considered in this study.^37^ Finally, we applied null modelling techniques to determine the drivers of microbial community structure and used niche breadth analysis to capture the relationship between microbes and continuous variables by parameterizing, as a model parameter, all possible gut environments in the Pakistani populace that these microbes were observed in. Thus, we provide a comprehensive analysis of antimicrobial resistance and the factors that influence microbial community structure in a Pakistani cohort.

## Materials and Methods

A detailed description of the experimental design, participant recruitment and screening, sample collection and sequencing analysis is included in our initial characterization work.^34^ Briefly, 32 ‘Gut’ and 28 ‘Oral’ samples were analyzed in total obtained from young healthy adults with age range (18-40) and BMI (18-25 kg/m^2^) with summary statistics are given in Table 1. The individuals recruited represent the major ethnic groups^♦^ across the country including Punjabi, Pathan, Saraiki, Kashmiri, Sindhi, and Balochi and belong to major geographic regions including Punjab, Khyber Pakhtunkhwa (KPK), Baluchistan, Sindh, Azad Jammu and Kashmir (AJK) and Federal Capital Islamabad. Out of the total of 32 individuals, 7 were reported to consume antibiotics three months or more before sampling. The study protocol was approved by the COMSATS ethics review board. All study participants provided written informed consent and data analysis was performed anonymously. The study was performed under a Human Subjects Protocol provided by an internal review board (E&I Review Services, IRB Study #13044, 05/10/2013)

**Table 1.** Summary statistics of the samples analyzed. A total of 60 samples were assigned to four major groups: Gut Male, Gut Female, Oral Male and Oral Female. Summary statistics, including median and Interquartile Range (IQR) of continuous variables such as *Age* and *BMI* is given.

|  | Groups | Total Samples (n) | Ethnic Groups (n) | Age (Years) |  | BMI (Kg/m2) |  | Smoking (n) |  | Antibiotics Usage (Past 3 months) (n) |  | Fresh Fruits (n) |  | Junk food (n) |  | Sources of drinking water (n) |
| --- | --- | --- | --- | --- | --- | --- | --- | --- | --- | --- | --- | --- | --- | --- | --- | --- |
|  |  |  |  | 18-40 |  | 18-25 |  | Yes | No | Yes | No | Yes | No | Yes | No |  |
| 1 | Gut Male | 15 | Balochi:1<br>Kashmiri:1<br>Pathan:8<br>Punjabi:4<br>Saraiki:1 | 23 | 19.50-25.50 | 22.4 | 20.50-24.40 | 5 | 10 | 2 | 13 | 4 | 11 | 13 | 2 | Bottled: 2<br>Filtered: 4<br>Mineral: 3<br>Tap: 6 |
| 2 | Gut Female | 17 | Kashmiri:2<br>Pathan:2<br>Punjabi:10<br>Saraiki:2<br>Sindhi:1 | 24 | 20-24 | 22.3 | 21.3-23.9 | 3 | 14 | 5 | 12 | 7 | 10 | 9 | 8 | Bottled:1<br>Filtered:12<br>Mineral:2<br>Tap:2 |
| 3 | Oral Male | 13 | Balochi:1<br>Kashmiri:1<br>Pathan:8<br>Punjabi:2<br>Saraiki:1 | 23 | 19-25 | 21 | 20.10-24.70 | 4 | 9 | 2 | 11 | 3 | 10 | 11 | 2 | Bottled: 2<br>Filtered: 3<br>Mineral: 2<br>Tap: 6 |
| 4 | Oral Female | 15 | Kashmiri:2<br>Pathan:2<br>Punjabi:9<br>Saraiki:2 | 24 | 22-24.50 | 21.9 | 21.10-23.85 | 3 | 12 | 4 | 11 | 6 | 9 | 7 | 8 | Bottled: 1<br>Filtered: 10<br>Mineral: 2<br>Tap: 2 |

### Sample Collection, DNA extraction and 16S rRNA amplicon sequencing

For sample collection, uBiome explorer kits were used and the participants self-collected stool and oral samples at home with the details of sampling and DNA extraction protocols given in ^34^. The samples were amplified for V4 region of 16S rRNA gene and were sequenced on the NextSeq 500 platform producing 2 x 150bp paired-end reads.

### Bioinformatics

The resulting 16S rRNA gene sequences (9,961,420) for n=60 samples were processed with the open-source bioinformatics pipeline QIIME2.^38^ Initially, the paired-end sequences were demultiplexed and quality trimmed using a PHRED quality score of 20. Deblur algorithm within QIIME2 ^39^ was then employed to recover Amplicon Sequence Variants (ASVs) which were then aligned to the reference alignment database SILVA SSU Ref NR release v138.^40^ Within QIIME2, q2-alignment method, MAFFT ^41^ was used to create multi-sequence alignment of ASVs, afterwards, a mask is applied to remove phylogenetically ambiguous alignments to obtain rooted phylogenetic tree using FastTree ^42^ within q2-phylogeny framework. After the bioinformatics steps, we obtained 2293 ASVs for n=60 samples with summary statistics of reads aligned per sample to these ASVs as follows: [1^st^ Quartile: 58,459, median: 124,589, mean: 127,625; 3^rd^ Quartile: 186,816; max: 282,077]. PICRUSt2 algorithm as a QIIME2 plugin ^35^ was used on the ASVs to predict the functional abundance of microbial communities. Although the prediction process is highly dependent on the number of pathways available for the reference genomes, with PICRUSt2, by virtue of a comprehensive reference database (∼20,000 genomes), only 2 out of 2293 ASVs were dropped out, not adhering to weighted Nearest Sequenced Taxon Index (NSTI) scores of 2.0 (average branch length that separates each ASV from a reference weighted by the abundance of ASVs), and as a result, this high alignment increases our confidence on the prediction quality.

### Statistical Analysis

R scripts were used for statistical analysis available at http://userweb.eng.gla.ac.uk/umer.ijaz/bioinformatics/ecological.html and some as part of R’s microbiomeSeq package http://www.github.com/umerijaz/microbiomeSeq. Before statistical analyses, we applied additional quality control measures such as the removal of likely contaminants (sequences identified as mitochondria or chloroplasts for example) and removed samples with < 5000 reads. The summary statistics of the final metadata are given in Table 1. Statistical analysis was performed using R software (R Core Team, 2022).^43^ All figures in this study were generated using R’s ggplot2 package.^44^

#### Microbial Diversity Analyses

The Vegan package ^45^ was used to calculate alpha and beta-diversity measures. Full details are described in the Supplementary Material.

#### Contribution of Antimicrobial Resistance Genes to Microbial Diversity

First, we defined AMR genes as those found from KEGG signatures of antimicrobial resistance genes (https://www.genome.jp/kegg/annotation/br01600.html), this gave a total of 90 KOs related to AMR (Supplementary Table 1). Out of the full list of 90 KOs, 52 were observed in our dataset. To see how these AMR genes vary within a cohort (i.e., gut versus oral samples, organized by *Antibiotics Usage Status* and *Smoking Status*), we calculated how much these genes contributed to beta diversity patterns observed in the dataset. For this purpose, we employed the *Bray-Curtis* (BC) dissimilarity as a metric of community dissimilarity, and is defined as 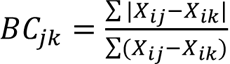, where *BC* is the Bray-Curtis dissimilarity between communities *j* and *k* and *X* is the relative abundance of the KEGG orthologs *i*. Since BC is a scaled summation of abundance differences between two communities, we can easily partition BC dissimilarity between two samples attributable to the AMR KOs (Supplementary Table 1). To obtain the contribution of AMR KOs, for two sample, we calculate the BC twice, first calculating the summation in the numerator of the BC expression but using the subset of KOs (BC_subset) relevant to AMR, and once with all the total KOs (BC_all). There is no change in denominator. Dividing BC_subset by BC_all then reports the fraction of beta diversity attributed to AMR genes. For this purpose, we employed R’s otuSummary package.^46^ To see if the Bray-Curtis contribution was significant between groups, we used Tukey Honest Significance Difference (HSD) test from R’s Stats package.

#### Interaction between Antimicrobial Resistance (AMR) KEGG Orthologs (Kos) and Study Participant Variables

To find the relationship between microbial communities/AMR KOs and sources of variation (*Age*, *BMI*, *Smoking Status*, *Antibiotics Usage Status*, *Fresh Fruits Status*, *Junk Food Status*, and *Sources of Drinking Water*), we have used Generalised Linear Latent Variable Model (GLLVM) ^37^ which extends the basic generalized linear model that regresses the mean abundances of microbes/AMR KOs against all sources of variation, even those that are not directly observed, as confounding latent variables. Further details are described in the Supplementary Material.

#### Differential Taxa

Since much of the data was paired, i.e., same individuals provided two samples, one for gut, and one for oral samples, we have used a specialised cluster association test ^47^ utilising R’s miLineage package (https://tangzheng1.github.io/tanglab/software.html) with this test referred to as QCAT-C test using the QCAT_GEE. Cluster() function (with default values) from the package. The test is robust to deduce complex correlations that exist among microbes due to paired nature of samples. Additionally, the QCAT-C test is a two-part test where it fits separate models to microbes that are excessively zero, and those that are not, referred to as positive microbes, based on the taxonomic tree to localize the covariate-associated lineages (*Gut Male*, *Oral Male*, *Gut Female*, and *Oral Female*). As a result, the differential abundance analysis of microbes gives better estimates and reduces Type 1 errors. To visualise the differentially abundant taxa at different taxonomic ranks, we have used Total Sum Scaling followed by a Centralized Log Ratio (TSS+CLR) transformation on the raw abundance values.

#### Microbial Niche Width

To assess microbial niche width, we have used R’s MicroNiche package ^48^ with the goal of determining positively/negatively associated microbial species with *BMI* and *Age* by explicitly incorporating the possible set of environments as a parameter (something that simple correlation analyses excludes). We have run the algorithm for *Age* and *BMI* separately for all possible combination of *Ethnicity x Antibiotics Usage Status*, as well as *Ethnicity x Smoking Usage Status*, considering these as possible set of environments a microbe is exposed to, for four different cohorts, i.e., *Gut Male*, *Oral Male*, *Gut Female*, and *Oral Female*. Further details of the implementation are described in the Supplementary Material.

#### Complexity Stability Analysis

To understand complexity-stability relationship of gut and oral cohort, we have followed Yonatan et al. (2022) computational framework for estimating the complexity of microbial ecosystems.^49^ Typically, an ecological system with *n* interacting species can be modelled with a Generalised Lotka Volterra model of the form 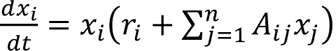, where *x_i_* is the abundance of species *i*, *r_i_*, is the intrinsic growth rate of species *i*, and *A_ij_* is the interaction coefficient, i.e., the effect of species *j* on species *i*. Traditionally, this matrix was obtained through network inference approaches that usually do not capture the causal relationships between interacting species in *A_ij_*. Once the interaction matrix is obtained, the stability-complexity relationship of the ecosystem can be derived as a curve that satisfies 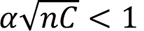 (also called May’s stability criteria ^50^ where *α*^2^ and *C* are the variance and density (“connectance”) of the non-zero off-diagonal elements of *A_ij_*. The effective connectance *D*^2^ is then inferred (where *D*^2^ ∝ *α*^2^*C*) without the need to infer a co-occurrence relationship, which is obtained by the slope of regression fitted to the dissimilarity/overlap plot to the 25% overlap values for the paired-wise dissimilarity/overlap values for *N* samples in a given category (oral or gut) from a total of *N*(*N* − 1)/2 paired-wise values. To calculate the dissimilarity between samples, we have used Spearman correlation, whereas to calculate the effective number of species, we used richness as an exponential of Shannon entropy.

#### Null-modelling Analyses

To understand the ecological mechanisms that are driving microbial community assembly in the gut and oral communities, we used various null modelling tools. Briefly, we used the Nearest Taxon Index (NTI) to determine whether the community was structured due to strong *environmental pressure* based on local clustering in phylogenetic tree. NTI is typically preferred for microbial datasets due to presence of significant phylogenetic signals across short phylogenetic distances.^51^ The implementation of this method is outlined in the Supplementary Material. We further used the normalised stochasticity ratio (NST) to determine the ratio of stochasticity in the communities.^29^ Taxa-richness constraints of proportional-proportional (P-P) and proportional-fixed (P-F) were applied for each metric. We have also used permutational multivariate ANOVA (also referred to as PANOVA) to obtain significance for NST between different cohorts. We then applied Quantitative Process Estimates (QPE) to quantify the contribution of specific assembly processes. This is based on an ecological framework that breaks down assembly processes into *Variable* or *Homogeneous selection*, *Dispersal limitation* or *Homogenising dispersal* and *Undominated* (neither dispersal nor selection processes dominate).^29^ *Variable selection* gives rise to high compositional differences in community structure due to different environmental/host conditions, whilst *Homogenous selection* occurs when these conditions result in consistent pressure. Dispersal processes refer to the movement of microbes throughout space, whether they are absent (*Dispersal limitation*) or present with high rates (*Homogenising dispersal*), resulting in homogenisation. Further details can be found in the Supplementary Material.

## Results

### Diversity patterns differ across gender and sample types

We have used several measures of alpha diversity including those derived from Hill numbers (parameterized with q [diversity index]), with q = 0 (Richness), q = 1 (Shannon Entropy), and q = 2 (Simpson) with increasing emphasis on abundant species. Generally, species richness (total number of expected species) in the gut samples was higher than the oral samples, and the gut of female participants showed higher richness than those of males (Figure 1A). These differences were statistically significant for Pielou’s evenness (balance in species numbers) and Simpson (putting more emphasis on abundant species) indices. When considering beta diversity through abundance counts alone, we can see two distinct clusters of oral and gut samples, with ∼28% variance explained by sample types using PERMANOVA (Figure 1B). Whilst there was a marginal difference (p=0.059) in microbial composition between males and females, accounting for 2% variation, these differences became significant (p<0.001) when phylogeny was considered (weighted UniFrac). This results in two distinct gut microbial profiles from males and females, retaining an overall ∼26% variation between the gut and oral samples (Figure 1B).

**Figure 1.**
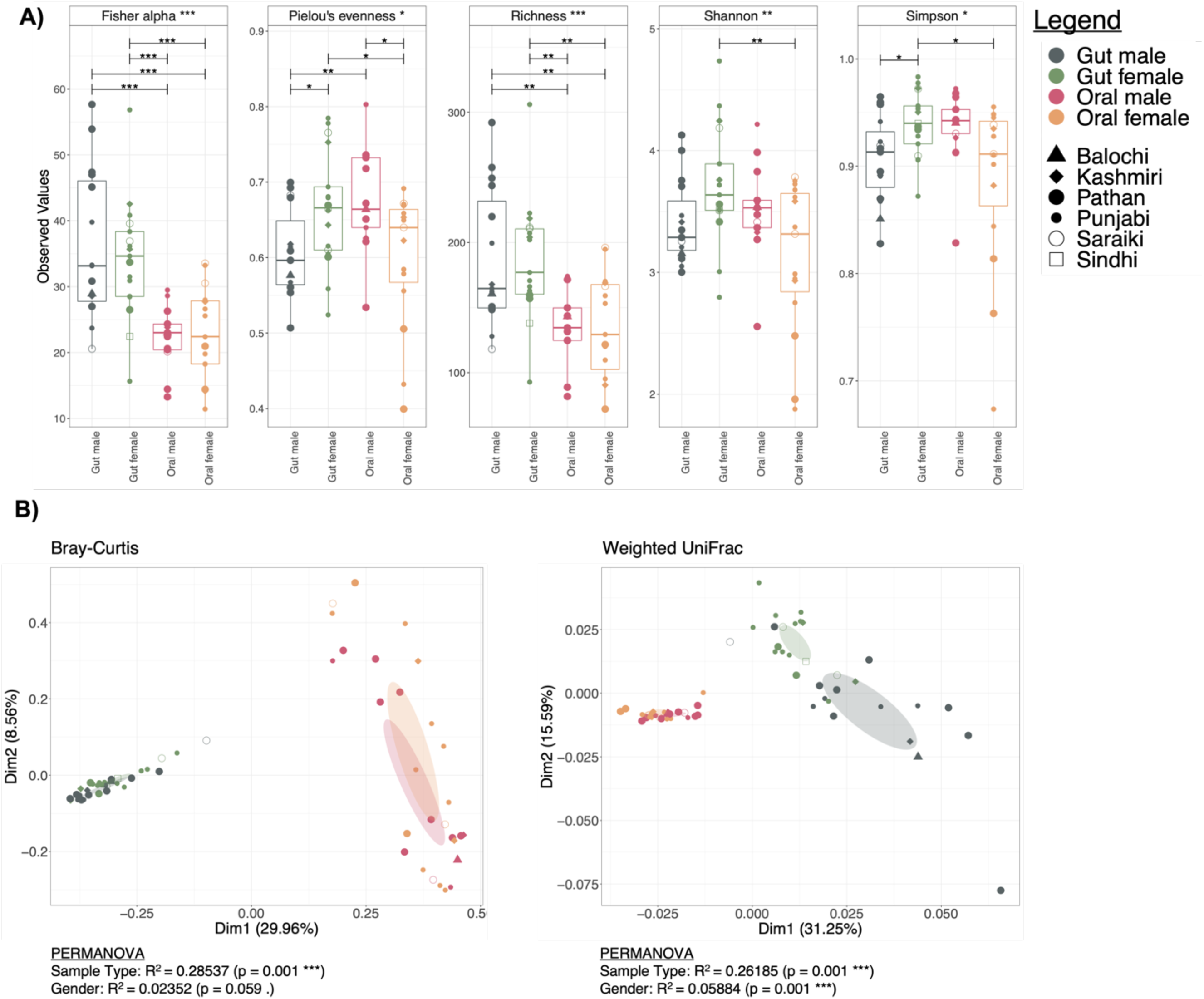
Diversity estimates. **A)** Commonly used alpha diversity indices with overall significance considering all four cohorts (gut male, gut female, oral male, and oral female) represented in the strip legends whilst pair-wise differences if significant are represented by lines connecting two categories. **B)** Principal coordinate analysis (PCoA) using different several dissimilarity indices (Bray-Curtis and Weighted UniFrac) where ellipses were drawn using 95% confidence intervals based on the standard error of ordination points for a given category. Beneath each figure are the R^2^ values (along with p-values if significant) calculated from PERMANOVA.

### Inter-sample variability in AMR genes is attributed to both antibiotic usage and gender

Next, we determined how AMR genes may have contributed to the observed microbial profiles. For this purpose, we recovered AMR KEGG KOs using PICRUSt2, and determined how these KOs contributed to Bray-Curtis dissimilarity. Signatures of AMR composition were similar and much lower in contribution than any other cohort for oral samples where antibiotics were not used (Figure 2). However, the contributions were highly variable within sample groups, though we observed less variability in oral samples as compared to gut samples. For example, the contribution of AMR genes ranged from 0.05% to 0.3% in the gut of females who declared they had not taken antibiotics, as compared to 0.02% to 0.1% for the female oral samples. In terms of *Smoking Status*, a similar trend in AMR composition was found for most sample types, except the female samples which were lower in contribution, however, these results are inconclusive due to the smaller sample size of females who smoked (n = 3).

**Figure 2.**
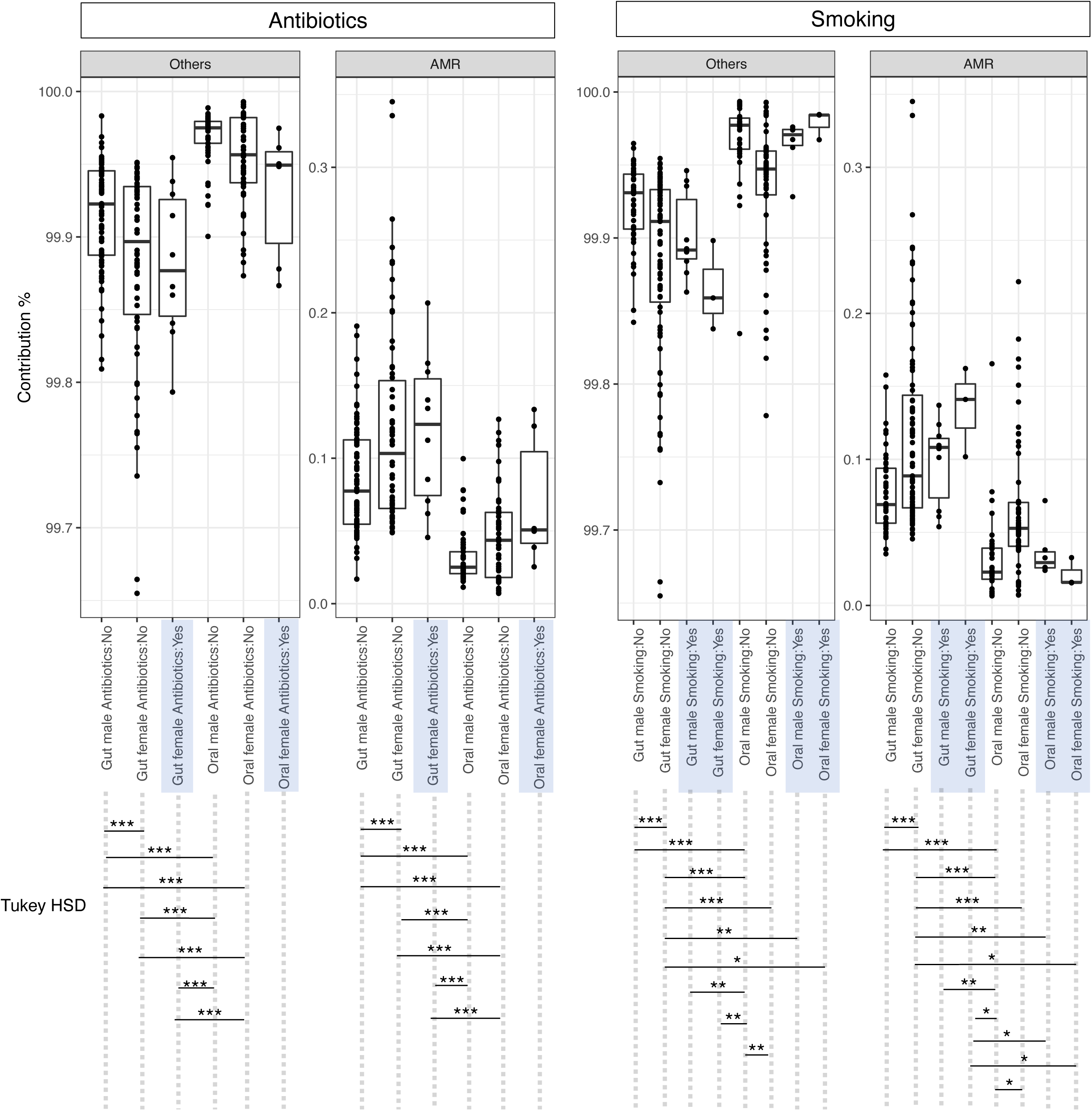
Bray-Curtis contribution of AMR genes to the overall beta diversity within the categories. The plot shows *N*(*N* − 1)/2 pair-wise differences between samples for each category with left set of panels showing organization by antibiotic status whilst right set of panels show organization by *Smoking Status*, respectively. “Others” represent the contribution of KOs that are not considered as AMR genes. Higher contributions represent higher inter-sample variability in terms of AMR genes.

### Sources of variations implicated in the prevalence of antimicrobial resistance genes

We then analyzed the abundance of KOs associated with AMRs against all sources of variation using generalized linear latent variable models (GLLVM) to find the covariates that on average caused a substantial change in the abundance of AMR genes. The variables tested were as follows: participant’s *Age*, *BMI*, *Gender*, *Sample type* (gut and oral), *Antibiotics Usage Status*, and *Dietary Habits* variables (Junk Food and Fresh Fruits), *Source of drinking water* (tap, mineral or filtered) and *Ethnic groups* (Kashmiri, Pathan, Punjabi, Saraiki, and Sindhi). Interestingly, in the participants that declared antibiotic use status “Yes” (7 gut and 6 oral samples), only 5 AMR genes (out of 52) were found to have a strong positive association with antibiotic use (Figure 3; K19217, K19322, K19276, K17881, K19100). These AMR KOs belong to beta-lactamase and aminoglycoside drug groups and are associated with pathogens that are classified as serious threats by CDC, including multidrug-resistant *Acinetobacter* and multidrug-resistant *Pseudomonas aeruginosa*. Out of 52 AMR KOs, 11 were found to be increased with increasing *BMI*.

**Figure 3.**
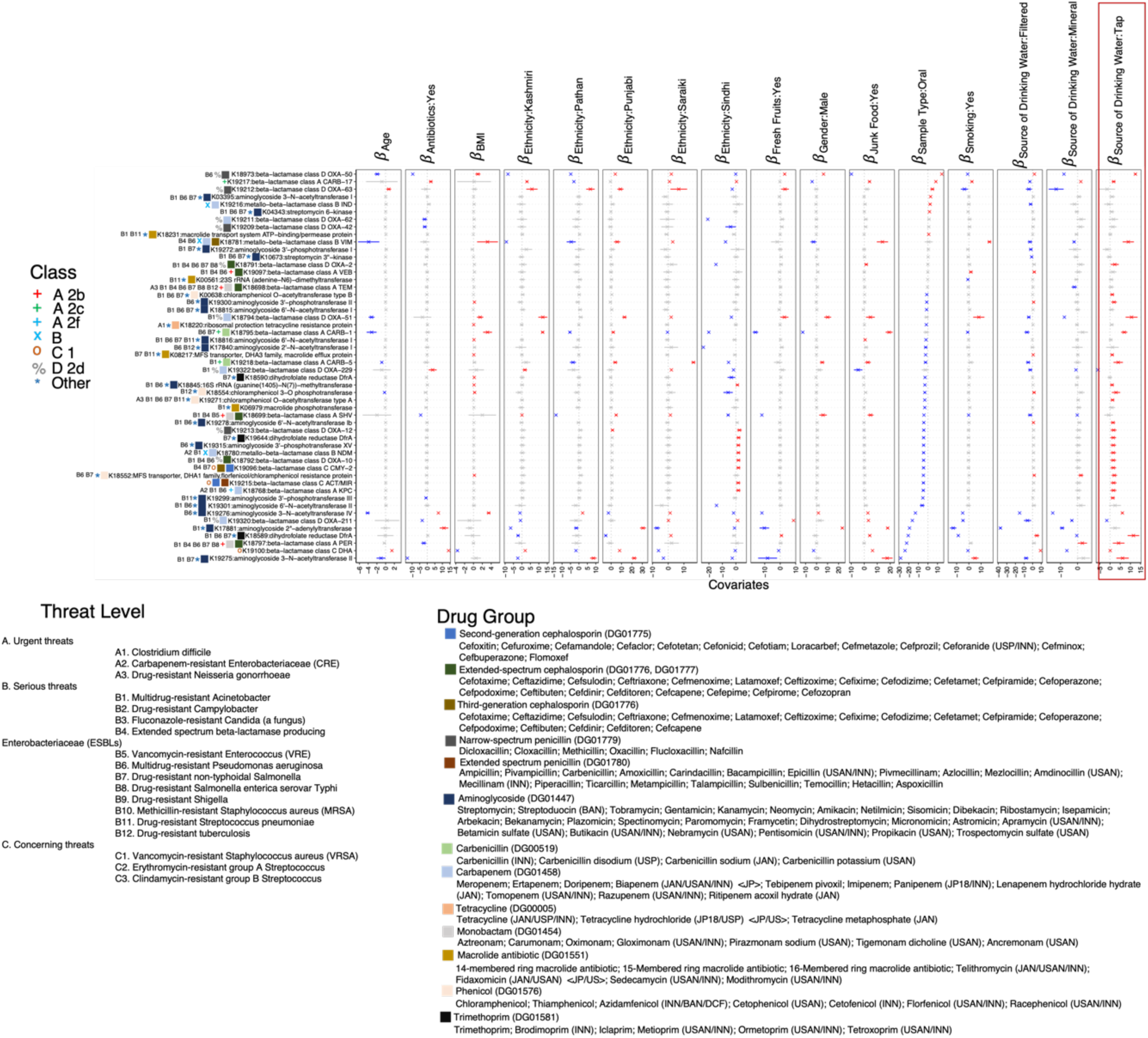
*β* −coefficients returned from GLLVM procedure for covariates considered in this study by considering 52 out of 90 reference AMR KEGG Orthologs (KOs) detected in this study using PICRUSt2 procedure. Those coefficients which are positively associated with the abundance of a particular AMR KO are represented in red colour whilst those that are negatively associated are represented with blue colour, respectively. Where the 95% confidence interval of the *β* −coefficients cross the 0 boundary, the coefficients are insignificant and are represented by grey color. Additional annotations comprise of threat levels, classes, and drug groups these AMR KOs are categorized under based on KEGG signatures of antimicrobial resistance genes (https://www.genome.jp/kegg/annotation/br01600.html; Supplementary Table 1). The red box is drawn around the *Drinking Water: Tap* to highlight it as a covariate with most numbers of positively associated *β* −coefficients.

Most of these KOs belong to beta-lactamases including narrow-spectrum penicillin and extended-spectrum cephalosporin drug groups. Drinking water sources with tap water showed 29 positively associated KOs as compared to 3 with bottled mineral water. Most of these AMR KOs were associated with serious threat pathogens (B6, B7, B1) and some with urgent threats (A2 and A3). Pathogens in urgent threats included; A2: Carbapenem-resistant *Enterobacteriaceae* (CRE) and A3: Drug-resistant *Neisseria gonorrhoeae*. In terms of sample types, there were more positive associations of AMR with Gut samples (33/52) as compared to oral samples (7/52), and most of them belong to beta-lactamases and aminoglycoside drug groups. For dietary intake, especially those consuming high carbohydrate and high fat that is *Junk food*, 14 AMR KOs having positive beta coefficients. Individuals consuming *Fresh fruits* showed only 5 AMR KOs with positive beta coefficients. One of the variables was *Ethnicity*, fewer AMR KOs with positive beta coefficients were found in Pathan 3/52, as compared to Punjabi, having 11/52 positive associations of AMR. There was less variation observed considering gender type, with only 5/52 positive association with males and 3/52 association with females and each of these AMR KOs belonged to beta-lactamases (Figure 3).

We also found that many of the AMR KOs detected were co-occurring (Supplementary Figure 2). For example, we found K19100 (beta−lactamase class C DHA), K19322 (beta−lactamase class D OXA−229) and K18768 (beta-lactamase class A KPC) each showed strong positive co-occurrence with K19315 (aminoglycoside 3’−phosphotransferase-XV), K18780 (metallo−beta-lactamase class B NDM) and K19096 (beta−lactamase class C CMY−2). While, K18795 (beta-lactamase class A CARB-1 [Carbenicillin]), K18973 (beta-lactamase class D OXA-50 [Narrow spectrum Penicillin]), and K18781 (metallo-beta-lactamase class B VIM [Third-generation Cephalosporin]) showed inverse co-occurrence with K18781 (metallo−beta-lactamase class [Extended spectrum Cephalosporins and Carbapenem]), K19320 (beta−lactamase class D OXA [Carbapenem]), K18792 (beta−lactamase class D OXA−10 [Extended Spectrum Cephalosporin]), K18794 (beta−lactamase class D OXA−51 [Carbapenem]), K19218 (beta−lactamase class A CARB–5 [Carbenicillin]) and K18797 (beta−lactamase class A [Extended spectrum Cephalosporin]).

### Key microbes associated with sources of variation

Using the variables outlined above (*Age, BMI, Antibiotic Usage Status, Smoking, Ethnic groups, Dietary Habits,* and *Source of drinking water)*, we again applied GLLVM, in this case, to find the microbial taxa that were strongly associated with the variables that showed strong positive associations with AMR KOs. The taxa that were positively associated with *drinking tap water* were *Brachyspira*, *Phascolarctobacterium, Butyrivibrio, Succinivibrio*, *Gastroanaerophilales*, *Neisseria*, *Strepobacillus*, *Cardiobacetrium*, and *Tannerella* (Figure 4). *Brachyspira* taxa showed a strong positive association with the male gender and, additionally, the use of antibiotics. *Phascolarctobacterium* also showed a strong positive association with *Antibiotic use*. While we observed reduced *Megamonas* abundance with *antibiotic use*, and a similar negative association with increasing *Age*. In contrast, we observed a strong positive association of *Megamonas* with *smoking*. *Lachnoanaerbaculum*, *Bergeyella*, *Tannerella*, *Cardiobacterium*, *Stomatobaculum*, *Streptobacillus*, *Pseudopropionibacterium*, and *Selenomonas* taxa groups were all positively associated with oral samples (Figure 4). Additionally, we observed differences in the microbial taxa with *Ethnicity*, with Saraki individuals showed a positive association with *Brachyspira* and other taxa which were negatively associated with Punjabi, Pathan and Kashmiri. Our data also highlighted certain microbial taxa co-occurrences (Supplementary Figure 3). For example, *Neisseria*, *Actinobacillus* and *Megasphaera* showed an inverse co-occurrence with other bacterial taxa including *Ruminococcus*, *Lachnospira* and *Bifidobacterium*. Moreover, *Blautia*, *Paraprevotella* and *Alistipes* showed a strong positive co-occurrence with *Bifidobacterium, Faecalibacterium*, and *Ruminococcus*.

**Figure 4.**
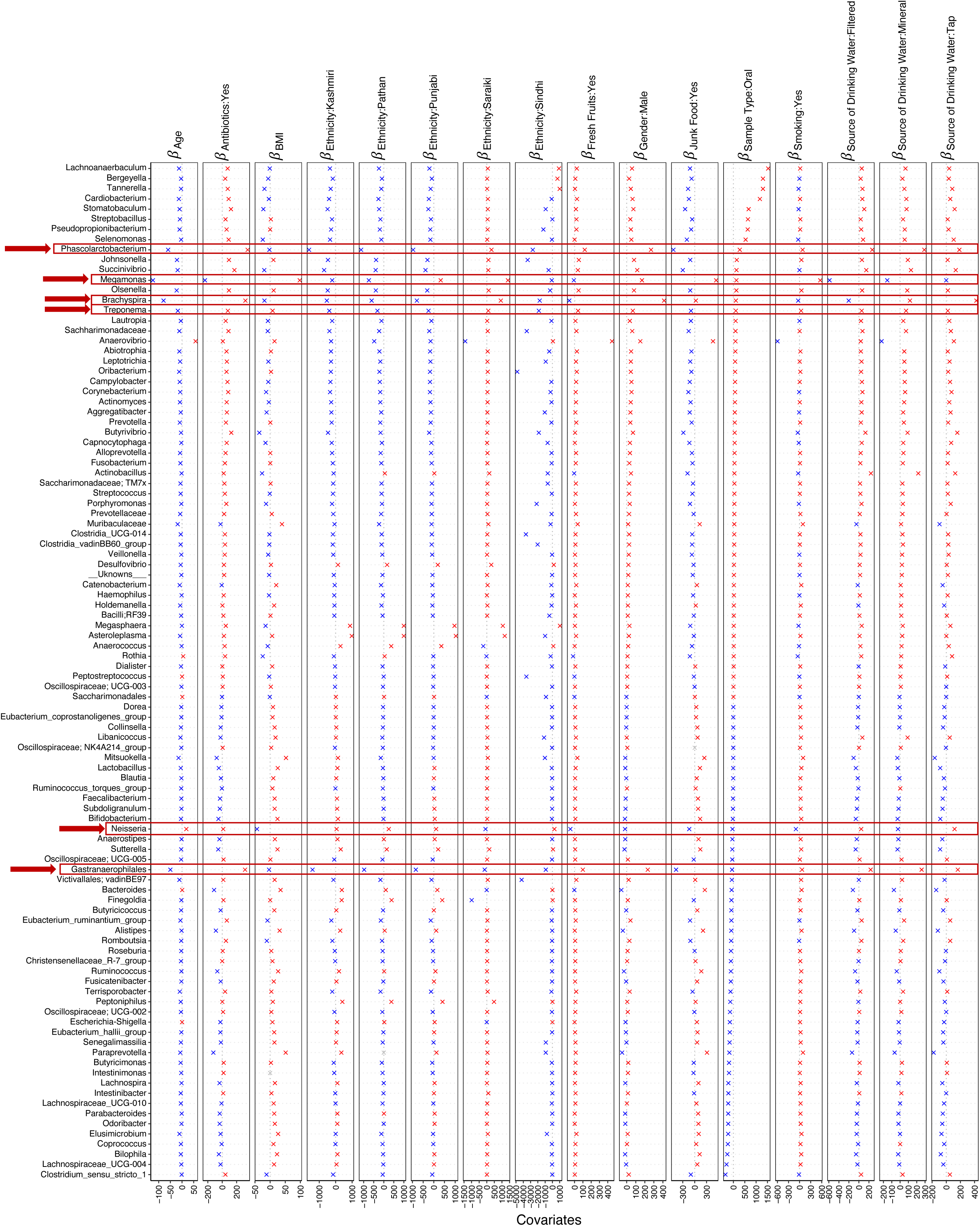
*β* −coefficients returned from GLLVM procedure for covariates considered in this study. Top 100 most abundant genera were considered, incorporating both continuous data (Age, BMI) as well as categorical labelling of samples. Those coefficients which are positively associated with the microbial abundance of a particular species are represented in red colour whilst those that are negatively associated are represented with blue colour, respectively. Since the collation of ASVs was performed at Genus level, all those ASVs that cannot be categorized based on taxonomy are collated under “__Unknowns__” category. Genera that standout in terms of significances with multiple covariates are highlighted with a red border.

### Microbial niche exploration reveals signature taxa for oral and gut communities associated with Age and BMI

To see how the environment shapes the distribution of taxa (i.e., what niche dictates which taxa should proliferate in a specific environment) we performed microbial niche analysis. This analysis was conducted using the numerical covariates (*Age and BMI*) and were further parameterized by *Ethnicity* and *Antibiotics Usage Status* or *Smoking Status.* In terms of *Age* and *BMI*, we observed three clusters, one for gut, and another one for oral samples as well as a third cluster, where genera change for the male gut with increasing *Age*, when the environments are a combination of *Ethnicity* and *Smoking Status* (Figure 5). Notably, the latter cluster shows only negative associations with *Megasphaera*, *Brachyspira*, *Phascolarctobacterium* and *Anaerococcus,* for example, while the former two clusters showed a mix of positive and negative associations with a wide range of taxa (Figure 5). Some of the genera that lie between the clusters for gut and oral samples simultaneously change in both types of samples (inverse or concomitant relationship between gut and oral) with respect to *Age* and *BMI*. These include *Treponema* “1” (inverse)*, Porphyromonas* “62” (inverse), *Streptococcus* “72” (inverse), *Corynebacterium* “41” (inverse), *Campylobacter* “5” (inverse), *Prevotella* “27” (concomitant), and *Alloprevotella* “34” (concomitant). With respect to the third cluster, there are genera from male gut samples that are negatively associated with *Age* (when the environments are parameterized in the model as a combination of *Ethnicity* and *Smoking Status*) and are not associated with any other cohort. These include *Finegoldia* “83”, *Elusimicrobium* “87”, *Peptoniphilus* “85”, *Libanicoccus* “86”, *Phascolarctobacterium* “81”, *Anaerococcus* “82”, *Tyzerella* “80”, and *Lachnospiraceae [Ruminococcus] gnavus group* “84”. Some genera were associated positively with *BMI* for male gut samples (including a combination of *Ethnicity* groups and *Antibiotic Usage Status*) and were not associated with any other cohort. These include *Romboutsia* “23”, *Mitsuokella* “3”, *Murdochiella* “8”, *Parabacteroides* “15”, *Desulfovibrio* “16**”,** *Clostridium_sensu_stricto_1* “21”, and *Oscillospirales;[Eubacterium] coprostanoligenes group* “17”. Considering *BMI* as a *property* (a continuous covariate observed in all possible environments) for gut male samples, there were a total of six environments observed in terms of *Ethnicity* and *Antibiotics Usage Status* (Supplementary Figure 3), the only microbial genus whose abundance decreased with increasing *BMI* was *Treponema*.

**Figure 5.**
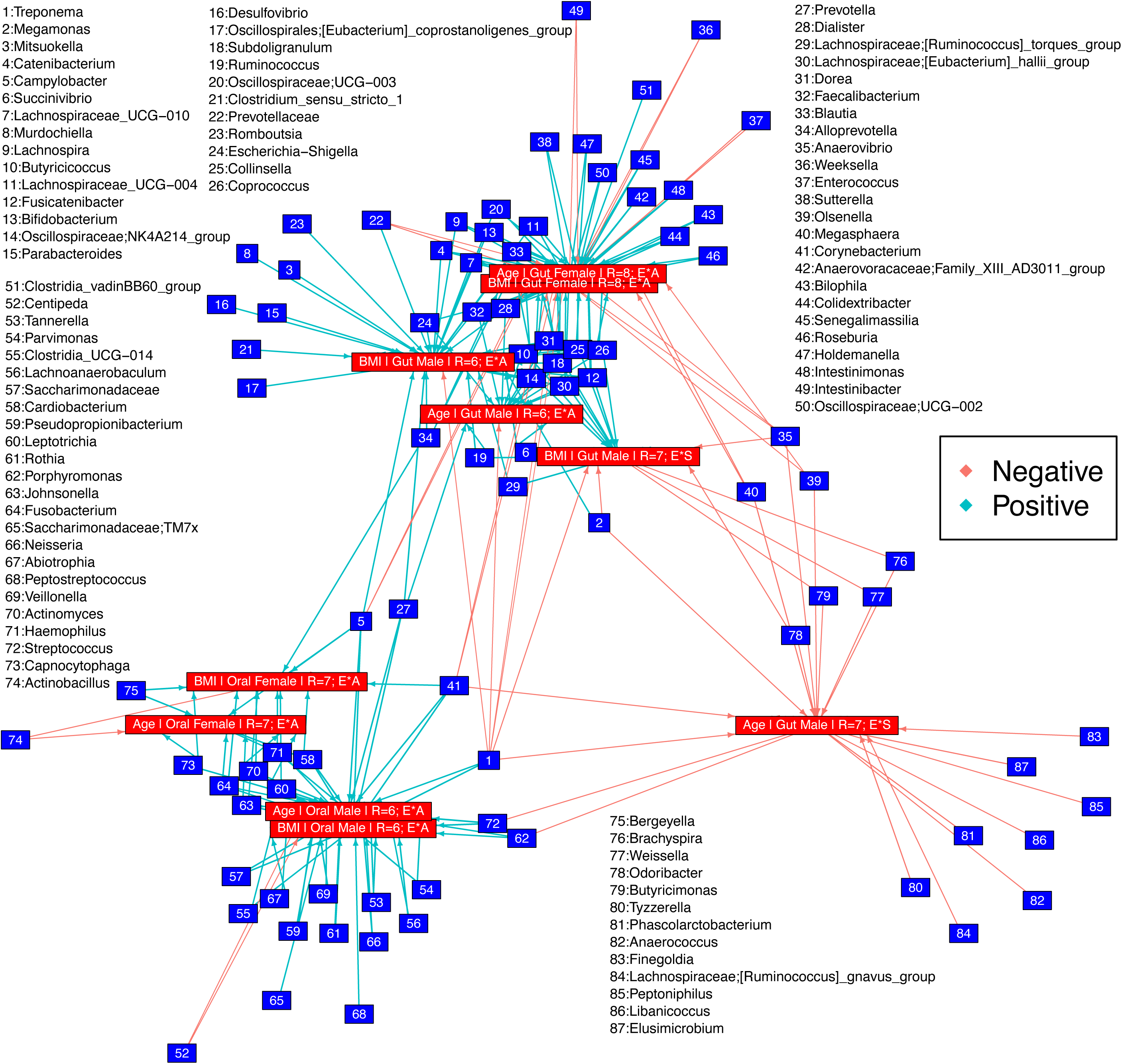
Relationships recovered after applying Hurlbert’s B_N_ to two environmental properties, *Age* and *BMI* on different sets of environments as a combination of *Ethnicity* and *Antibiotics Usage Status* (E*A) or *Ethnicity* and *Smoking Status* (E*S) with R representing the possible sets of environments. Genera tagged as “Positive” increase in abundance, whilst those tagged as “Negative” decrease in abundance in relationship to the environmental property considered as well as the target set of environments. Further details are given in Supplementary Figure 3.

### More environmental pressures on oral communities than gut communities

The above results highlighted that sample type was a distinct niche for specific taxa groups. Therefore, in this next section, we wanted to determine what could be driving the observed patterns in microbial communities. Using the nearest taxon index (NTI), we confirmed that for both gut and oral communities, a strong environmental pressure (NTI > +2) is exerted on these communities, that is dependent on the host environment they reside in (Figure 6A). Relatively, higher environmental pressure was observed on oral communities than gut communities. We applied normalized stochasticity ratio (NST) analysis (Figure 6B) to confirm this observation where, depending on the beta diversity measure and null model chosen, the oral communities were found to be more deterministic. Between genders, female oral communities are more deterministic than male oral communities with marginal significance in terms of PANOVA, however, the converse trend was observed for gut communities where the gut of females was the most stochastic amongst all the cohorts considered. The largest contribution to assembly process was *Dispersal Limitation* (Figure 6B). The emphasis here is on the deterministic measures of the assembly processes, where ∼12% of the oral male microbiome is explained by *Homogeneous Selection* suggesting similar environmental oral conditions. On the other hand, ∼10% of the female gut microbiome is explained by *Variable Selection* suggesting existence of multiple stable gut environment.

**Figure 6.**
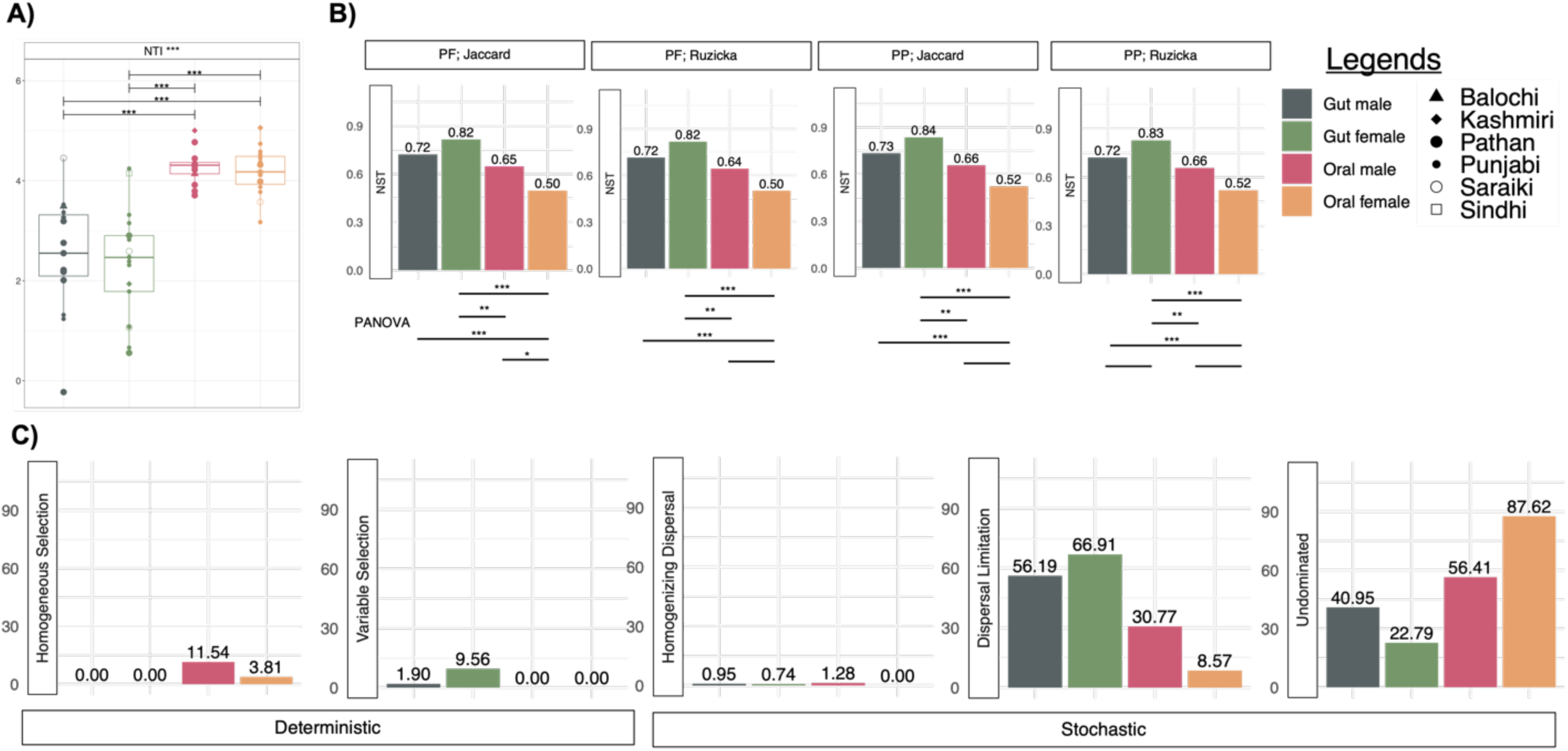
Estimates of microbial community assembly mechanisms structuring samples from four different cohorts (Gut Male, Gut Female, Oral Male, and Oral Female). **A)** phylogenetic alpha diversity measure, Nearest Taxon Index (NTI) to elucidate environmental pressures. **B)** Normalised stochasticity Ratio (NST) calculated as both incidence-based (presence/absence) Jaccard and abundance based Ruzicka metrics with PF and PP being the null modeling regime used. PANOVA connects pairwise categories where the estimates are statistically significant in terms of NST; **C)** Proportion of assembly processes returned from applying Quantitative Processing Estimate (QPE) for all four cohorts presented as both deterministic and stochastic measures.

### Oral microbiota is less stable as compared to the gut microbiota

We have performed complexity-stability analysis to find the complexity of microbial interactions within gut and oral communities. Higher effective connectance suggests that any local perturbation in the one or few species within oral microbiota can affect the entire community more substantially, making it less stable when compared with gut. We have found that *effective connectance* D^2^ value was higher for oral samples as compared to gut samples (Figure 7A), suggesting that oral communities are less stable when compared with gut microbial communities. D^2^ value is calculated using the square root of x-value which for oral communities was 1.83 (Figure 7B) compared to 0.70 for gut communities (Figure 7C).

**Figure 7.**
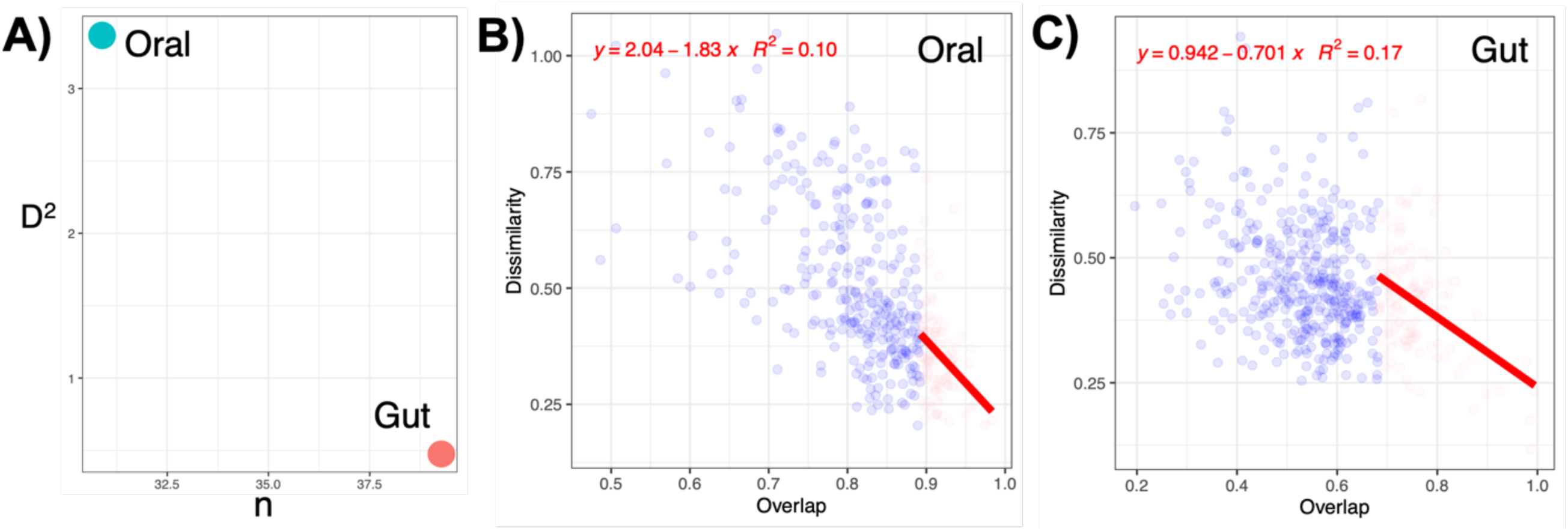
Complexity-stability relationship of oral and gut communities (**A**) where the effective connectance *D*^2^ is calculated based on fitting a linear regression to top dissimilarity/overlap values for pair-wise samples where the overlap values are in top 25% of communities, and then recovering *D*^2^ as the slope of fitted regression for oral (**B**) and gut (**C**) communities.

### The link between oral and gut microbial communities

Since most of the samples analyzed in this study were paired (coming from same subject), we have used an advanced paired differential analysis, i.e., QCAT-C (specialized cluster association test) analysis to analyse the microbial profiles between body sites (gut and oral). The method considers the taxonomy of all ASVs and returns the differential taxa at each taxonomic level (genus, family, order, class, phylum) with the differentially abundant taxa from these levels shown in Supplementary Figures 5-7. Interestingly, in those subjects (paired gut-oral samples from subjects are indicated using connected lines) that show specific taxa as outliers in the gut, these taxa were typically also highlighted as outliers in the subjects paired oral sample (e.g., at family-level: *Bifidiobacteriaceae, Butyricicoccaceae, Clostridiaceae, Microbacteriaceae, Neisseriaceae and Saccharimonadaceae* [Supplementary Figure 6] and at genus-level; *Alloprevotella, Campylobacter, Cardiobacterium, Bergeyella and Neisseria* [Supplementary Figure 7]). This differential analysis can also be used to determine the differentially abundant taxa between genders. For example, prominent taxa in the gut communities included *Elusimicrobium, Phascolarctobacterium and Succinivibrio*, which were in high abundance in males versus females (Supplementary Figure 7). In contrast, other prominent gut taxa, *Alistipes*, *Bacteroides* and *Bifidobacterium,* were high in female samples and low in male samples. Similarly, for oral communities, *Peptostreptococcus, Streptobacillus, Catonella, Cardiobacterium and Pseudopropionibacterium* were high in male samples and low in female samples. While *Centipeda, Monoglobus, Oscillibacter, Lachnospiraceae UCG 10, Desulfovibrio and Olsenella* were in high abundance than male oral communities (Supplementary Figure 7).

## Discussion

In the present study, we have provided an in-depth analysis of the gut and oral microbial communities within a Pakistani cohort. Pakistan is ranked third in the low and middle-income countries for frequent antibiotic consumption.^52^ Broadly, we highlighted numerous KEGG orthologs (KOs) associated with antimicrobial resistance (AMR) and linked prevalence to key variables such as *BMI* and *Drinking Tap Water.* We further linked these key variables with associations to specific taxa. Finally, we found differences in the estimated contribution of ecological assembly processes between the gut and oral communities of males and females. Overall, we have provided a comprehensive insight into the potential prevalence of AMR in Pakistani individuals and the factors driving the observed microbial community structure.

The gut and oral samples had a distinct microbial signature with gut samples showing greater diversity than the oral samples. The contribution of AMR genes to Bray-Curtis dissimilarity was higher in the gut samples, irrespective of antibiotic use, although values were highly variable, and the sampling space was low. Our results highlight that more AMR KOs increased in the gut as compared to the oral samples which is in agreement with the literature likely explained by differences in microbial host and mobile genetic element associations.^53^ In contrast, Carr et al. (2020) found that oral human microbiomes contained a higher abundance of ARGs than gut samples.^54^ However, our work is also consistent with the previous work highlighting the gut as a reservoir for antimicrobial resistance, although it did not consider oral communities.^55^ We used GLLVM to statistically determine the associations of AMR KOs with key variables. This revealed that more AMR KOs were associated with an increased *BMI* and notably were the highest in individuals who drank tap water. Around 80% of the Pakistani population is exposed to unsafe drinking water because of the scarcity of safe drinking water sources or lack of treatment facilities.^56^ It is therefore not surprising, that our analyses picked up *Drinking Tap water* as the main source of the spread of AMR.

The potential AMR KOs detected were associated with multi-drug resistant pathogens that belong to serious and urgent CDC threat levels. The majority of these were associated with cephalosporin resistance. This group of antibiotics is widely prescribed in Pakistani hospitals (67%).^12^ In addition, fluoroquinolones are also the most prescribed antibiotic groups in Pakistan.^57^ The presence of KOs in our study linked with carbapenem-resistant *Enterobacteriaceae*, *Acinetobacter baumannii* and drug-resistant *Neisseria gonorrhoeae* is also alarming. Bilal et al. (2021) reported that these bacteria show complete resistance towards a wide range of antibiotics in Pakistan including ceftriaxone, ciprofloxacin and cefepime.^11^ We thus speculate that poor treatment of hospital or sewage wastewater or agricultural run-off may be a source of contamination in the drinking water in Pakistan.^56, 58, 59^ In general, in LMICs, the most deadly and common pollutants in drinking water are of biological origin.^60^ For example, a study by McInnes et al. (2022) observed human waste as a primary source of AMR in Bangladesh.^61^

We further linked microbial taxa to individual variables (*Age*, *BMI*, *Smoking, Antibiotic Usage Status*, *Ethnic groups*, and *Source of drinking water*). Of these, *Megamonas* taxa had most significant associations: negatively associated with *Age*; and positively associated with *BMI* and *Smoking*. These are all in agreement with the literature.^62,63,64^ *Megamonas* abundance is linked with high caloric intake and restricted activity.^64^ Particularly, it is implicated in smokers and those who have a high BMI, which may likely be associated with endotoxemia and inflammatory phenotype of gut which is inherent in such subjects.^65^ More importantly, *Megamonas* species are involved in glucose fermentation into propionate and acetate ^66^ and these short-chain fatty acids (SCFAs) are beneficial for health.^67^ Studies have reported that SCFAs help in maintaining intestinal homeostasis ^68^ and can impact host immune response.^69^ Antibiotics alter the balance of SCFAs, and cause dysbiosis in the gut environment.^70^

We observed a strong positive association of antibiotics intake and drinking tap water with the *Brachyspira* taxa. There are a number of studies linking *Brachyspira* with diarrhea, and abdominal pain from faecal-oral transmission.^71, 72^ Studies have proposed that faecally contaminated water is an important route of *Brachyspira* transmission.^73^ Moreover, we identified a strong positive *Neisseria* genera with *Drinking Tap water*. This correlates to AMR KOs detected in our dataset corresponding to drug-resistant *Neisseria gonorrhoeae*, however, this link cannot be confirmed as amplicon sequencing cannot provide species or strain-level information. Nonetheless, even commensal *Neisseria* species have been linked to AMR transmission.^74^ Therefore, our findings merit further investigation of the drinking water microbiome in Pakistan for understanding AMR transmission and gut health.

Using microbial niche breadth approach, we highlighted that the gut and oral communities were distinct (when separated according to *Ag*e and *BMI*). Interestingly, the QCAT results showed that for some taxa, the subjects that were implicated as outliers in gut were also outliers in the oral samples suggesting that dysbiosis in the gut may also manifest in oral communities, e.g., literature suggests that the reported genera in our study including *Alloprevetolla* and *Campylobacter* are found in both gut and oral microbiota. The presence of *Campylobacter* in the oral microbiome is also linked to initiating inflammatory bowel disease (IBD).^75^ Indeed there are some recent studies where gut-oral axis is explored.^76, 77^ Along with that, we have also found AMR KOs with positive co-occurrences in both gut and oral communities. This warrants further investigation in a clinical setting as the use of oral communities as a proxy for gut communities and gastrointestinal system diseases would be more convenient.^78^

QCAT, GLLVM and niche breadth analyses highlight that certain bacterial genera including *Megamonas, Phascolarctobacterium* and *Succinivibrio* have high abundance in males, while *Bifidobacterium, Bacteroides* and *Ruminococcus* were more abundant in females. These results are in alignment with a Japanese study that reported higher levels of *Megamonas* in males and higher levels of *Bifidobacterium* in females.^79^ A Chinese study also reported a similar pattern with *Ruminococcus* as the most abundant genera in females.^80^ These studies suggest that in terms of microbial composition and most abundant genera, the Pakistani gut microbiome resembles the microbial profile within the Asian community. Increased abundance of *Proteobacteria* has been linked to gut dysbiosis^81^ and obesity^64^ and we have also reported an increased abundance of *Proteobacteria* in Pakistani males as compared to females in our previous study.^34^ This could be a cause for concern as a positive correlation of *Proteobacteria, Prevotella* and *Bacteroides* was associated with antibiotic resistance genes (ARGs) in a Pakistani cohort^82^ and we also observed a similar positive association of these genera including *Proteobacteria (Succinivibrio, Cardiobacterium, Actinobacillus* and *Aggregatibacter)* in Pakistani males.

Null modelling analyses further confirmed strong environmental pressure on the gut and oral microbial communities. This is not unexpected given that the host system places an additional constraint on microbial communities.^83, 84^ We found that dispersal limitation was a key assembly process in gut samples arising from the individual nature of each host, while oral communities showed weak selection and moderate drift (undominated) as the primary assembly process. Complexity stability analysis showed that the oral communities were less stable than the gut communities and thus may have more factors influencing microbial community assembly. Interestingly, we observed gender-based differences whereby female gut communities were influenced by variable selection, and male oral communities were influenced by homogenous selection. Dispersal limitation is reported as a primary assembly process in the human gut microbiome of US and Papa New Guinean populations as well.^85^ The authors also noted variable selection as a key process which was influenced by geographical location, however, the authors did not define study participants by gender. In the literature, analysis of the contribution of microbial assembly processes and how different covariates may impact the human microbiome is still relatively rare. Thus, our study provides valuable insight into the impact of the environment (gut, oral, male, and female) and lifestyle (*BMI, Age*, *Smoking*, and *Antibiotic Usage Status*).

Whilst we have based our AMR analyses on metabolic profiles predicted through PICRUSt2, it has been shown to perform well on human associated microbiome datasets.^35, 86, 87^ This is mainly due to a comprehensive reference database of genomes whose functions are already known (a 10-folds increase in the numbers since the previous release).^35^ Shotgun metagenomics of these samples would more accurately highlight AMR gene prevalence; however, the experimental cost and resources for processing and data analysis may be prohibitive for LMICs, and predictive modeling may be a viable monitoring option for AMR spread.

## Abbreviations

AMR: Antimicrobial Resistance
ARGs: Antibiotic Resistance Genes
ASVs: Amplicon Sequence Variants
BC: Bray-Curtis
BMI: Body Mass Index
CDC: Centers for Disease Control and Prevention
GLASS: Global Antimicrobial Resistance and Use Surveillance System
GLLVM: Generalised Linear Latent Variable Models
HGT: Horizontal Gene Transfer
IBD: Inflammatory Bowel Disease
IQR: Interquartile Range
KEGG: Kyoto Encyclopedia of Genes and Genomes
KOs: KEGG Orthologs
LMICs: Low and Middle-Income Countries
NIH: National Institute of Health
NST: Normalized Stochasticity Ratio
NTI: Nearest Taxon Index
QPE: Quantitative Process Estimates
SCFAs: Short Chain Fatty Acids

## Footnotes

♦ These are self-reported ethnicities identified by the participants at the time of sample collection, and were based on place of birth as opposed to where they were currently residing.

## Notes

**Grant Support:** The authors wish to thank the study participants for participating in the study and providing samples. The work is supported by EPSRC (EP/P029329/1 and EP/V030515/1) to U.Z.I., uBiome Academic Grant Program to A.N., and with contribution from Higher Education Commission, Pakistan Project No. 1-8/HEC/HRD/2022/1199 to M. B.

