## Supplementary Material for "Cross-sectional study of antimicrobial resistance and ecology in gastrointestinal and oral microbial communities of urban Pakistani adults"

^$^ Present address: Moderna, Inc., Cambridge MA, USA

**Supplementary Methods**

Study Area:

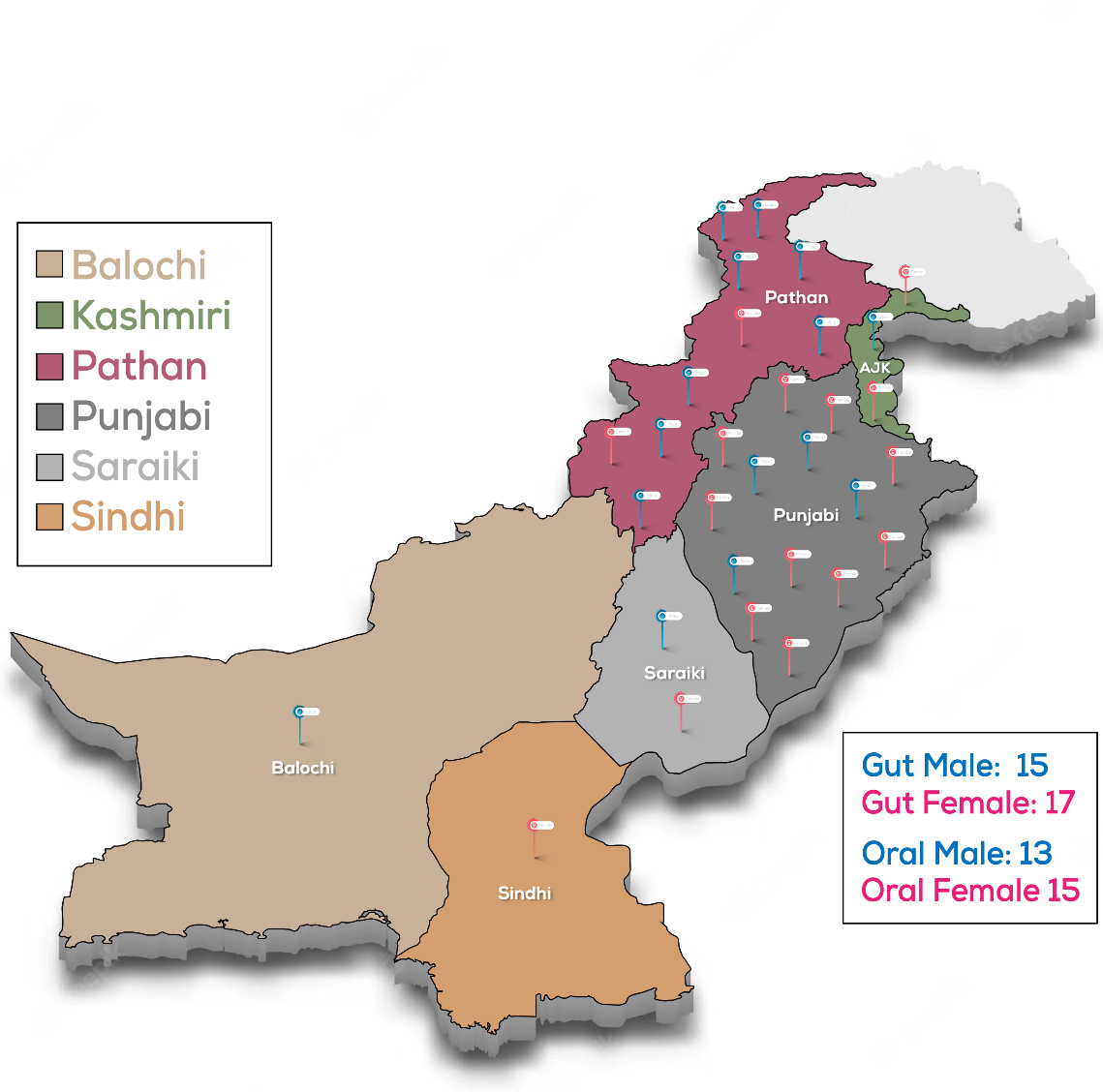

**Supplementary Figure 1.** Self-reported ethnicities of healthy individuals at the time of sample collection, which is primarily based on their place of birth (highlighted on the map). Majority of these samples were paired as the individuals provided both gut and oral samples.

**Statistical Analyses**

Prefiltering:

For statistical analysis, any samples with <5000 total reads were dropped, and also excluding contaminants based on taxonomy (Chloroplast, Mitochondria, and ASVs unassigned at Phylum level) as per typical recommendations given at <https://docs.qiime2.org/2022.8/tutorials/filtering/>.

Microbial Diversity Analyses:

For alpha diversity measures we have used: *Fisher’s alpha* – a parametric index of diversity that assumes the abundance of ASVs following the log series distribution; *Pilou eveness,* which compares the actual diversity values to the maximum possible diversity value, and is constrained between 0 and 1.0, where lower values will indicate more variation in abundance between different ASVs in the community; *rarefied richness* – the estimated number of species/features in a rarefied sample (to minimum library size); *Shannon entropy* – a commonly used index to measure balance within a community; and *Simpson index –* a measure of dominance that weighs towards the abundance of the most common ASVs and is less sensitive to rarer ASVs. Richness, Shannon entropy, and Simpson indices are parameterized versions of Hill numbers with q=0, q=1, and q=2, respectively. For beta-diversity, we have used principal Coordinate Analysis (PCoA) plots of ASVs using two different distance measures in Vegan’s cmdscale() function: *Bray-Curtis,* which is a distance metric that considers only ASV abundance counts; and *Weighted Unifrac,* which is a phylogenetic distance metric combining phylogenetic distance with relative abundances. Unifrac distances were calculated using the phyloseq package.^1^ Analysis of variance was performed using Vegan’s adonis() with distance matrices (Bray-Curtis/Weighted Unifrac) against sources of variation. This function referred to as PERMANOVA, fits linear models to distance matrices and used a permutation test with pseudo-F ratios and gives percentage variability in microbial community explained by a particular covariate as R^2^ value (if significant).

Interaction between Antimicrobial Resistance (AMR) Kegg Orthologs (Kos) and Study Participant Variables:

We have used Generalised Linear Latent Variable Model (GLLVM) ^2^ which extends the basic generalized linear model that regresses the mean abundances $\mu_{ij}$ (for $i$-th sample and $j$-th microbe/AMR KO) of individual microbes/AMR KOs against covariates $x_{i}$ by incorporating latent variables $u_{i}$ as $g\left( \mu_{ij} \right)=\eta_{ij}=\alpha_{i}+\beta_{0j}+\boldsymbol{x}_{i}^{T}\boldsymbol{\beta}_{j}+\boldsymbol{u}_{i}^{T}\boldsymbol{\theta}_{j}$, where $\boldsymbol{\beta}_{j}$ are the microbe/AMR KO specific coefficients associated with individual covariates. Further details are described in the Supplementary Material. (a 95% confidence interval of these whether positive or negative, and not crossing 0 gives directionality with the interpretation that an increase or decrease in that particular covariate causes an increase or decrease in the abundance of the microbe/AMR KO), and $\boldsymbol{\theta}_{j}$ are the corresponding coefficients associated with latent variable. $\beta_{0j}$ are microbes/AMR KOs specific intercepts, whilst $\alpha_{i}$ are optional sample effects which can either be chosen as fixed effects or random effects. To model the distribution of individual microbes and AMR KOs, we have used Negative Binomial distribution with an additional dispersion parameter, and using log() as a link function. Additionally, the approximation to the log-likelihood is done through Laplace approximation (LA) with final sets of parameters in glvmm() function being family = 'negative.binomial', method="LA", and starting.val='zero’ that seemed to fit well. This, we did for top 100 most abundant genera and all the AMR KOs observed in our datasets. In addition, the factor loadings $\boldsymbol{\theta}_{j}$ store correlations of microbes with the residual covariance matrix $\boldsymbol{\Sigma}=\boldsymbol{\Gamma}\boldsymbol{\Gamma}^{T}$ where $\boldsymbol{\Gamma}=[\theta_{1}\ldots\theta_{m}]$ for $m$ latent variables. This residual covariance matrix gave co-occurrence relationship between microbes and AMR KOs that is not explained by the observed covariates.

Microbial Niche Width:

Before implementing this approach, we filtered out genera by applying the limit of quantification (LOQ) method. LOQ fits a log abundance distribution and filters out taxa that fall below a decision threshold (1.65 x standard deviation of the fitted distribution), calculated from the distribution of microbes with 95% certainty that these microbes will fall within a null distribution where the mean microbial abundance is zero. With this pre-filtering step performed, we then calculated Hurlbert’s $B_{N}=\frac{1}{\sum_{i=1} \frac{p_{i}^{2}}{r_{i}}}$, where $p_{i}$ is the proportional abundance of a microbe in the $i$-th environment, and we have an additional $r_{i}$ proportional covariate data (BMI and Age) in the formula. The model yields a value between 0 and 1 for each microbe and corresponding covariate, indicating whether there is an inverse (~0) or a positive relationship (~1), with 0.5 indicating no relationship to the covariate. To obtain p-values, a null modelling procedure was considered by generating a random normal distribution of 999 possible Hurlbert’s $B_{N}$, and by tagging it as “negative” if its $B_{N}$ < 5^th^ Quantile, and “positive”, if its $B_{N}$ > 95^th^ Quantile, as per original author’s instructions. Finally, to visualise the results, we have used R’s igraph ^3^ to draw the network of relationships between the microbe and the observed covariates.

Null-modelling Analyses:

To calculate the Nearest Taxon Index (NTI) we have used R’s picante package to calculate it using mntd () and ses.mntd functions.^4^ We have used 999 randomisations using null.model=“richness” in the ses.mntd() function and have only considered taxa as either present or absent regardless of their relative abundance. Positive values of NTI indicate that species co-occur with more closely related species more frequently than expected by chance, with negative values suggesting otherwise. For NTI, values > + 2 indicate strong environmental pressure (determinism), and values < −2 indicate strong competition among species as the driver of community structure.

We applied the Quantitative Process Estimates (QPE) framework to quantify the contribution of different ecological assembly processes.^5^ The method uses deviation from the observed βMNTD (β-mean-nearest-taxon-distance) and the mean of the null distribution was evaluated using βNTI (β-nearest-taxon-index). When the observed value of βMNTD deviated significantly from the null expectation, the community is assembled by variable (βNTI >+2) or homogenous (βNTI < −2) selection processes. If the difference is not significant, the observed differences in phylogenetic composition are considered to be the result of dispersal mechanisms enabling ecological drift. These are differentiated using the abundance-based β_RC_ and a Bray-Curtis dissimilarity metric for beta diversity. Dispersal limitation contributes to the community assembly If the β_RCbray_ > + 0.95, homogenising dispersal will be contributing if the β_RCbray_ > + 0.95 and community turnover is due to undominated mechanisms if β_RCbray_ was between −0.95 and +0.95.

**Supplementary Results**

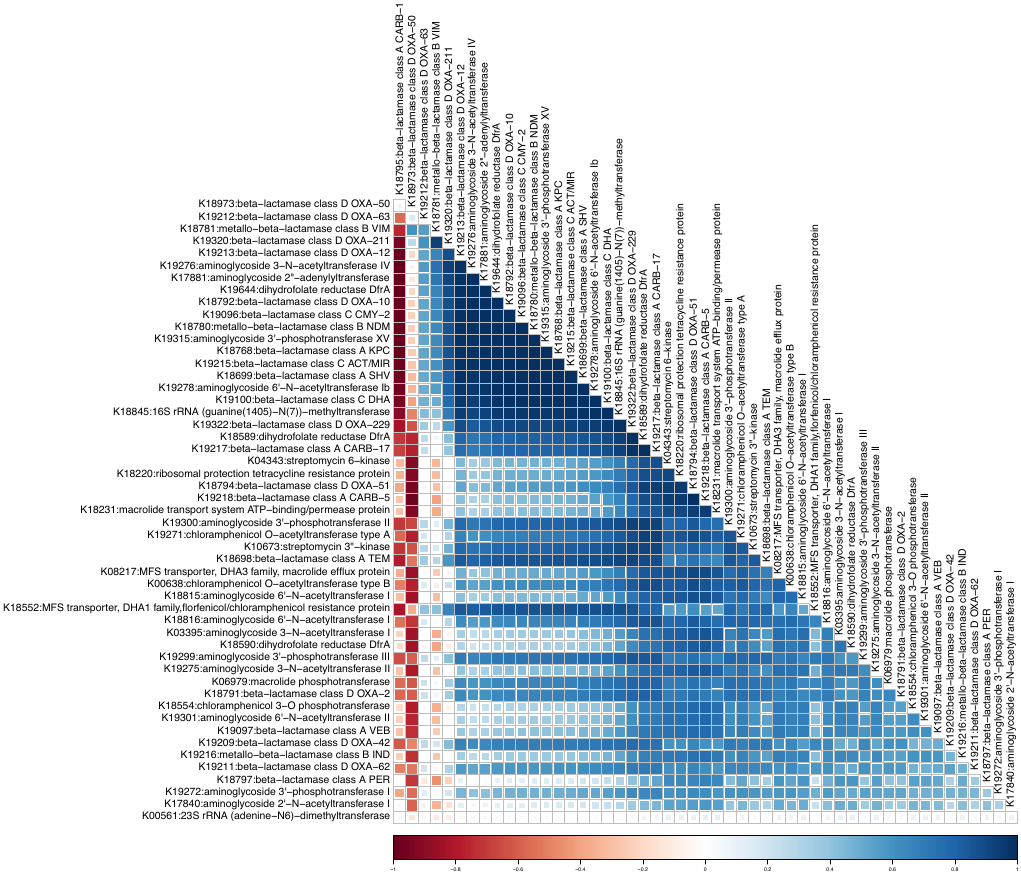

**Supplementary Figure 2**. Co-occurrence relationship between AMR KOs recovered from the residual covariance matrix $\boldsymbol{\Sigma}$ that are not explained by the observed covariates in the GLLVM model in Figure 5. Here, blue represent the positive correlation (KO of AMR1 increasing in abundance leads KO of AMR2 increasing in abundance), and red represent the negative relationship (KO of AMR1 increasing in abundance leads to KO of AMR2 decreasing in abundance).

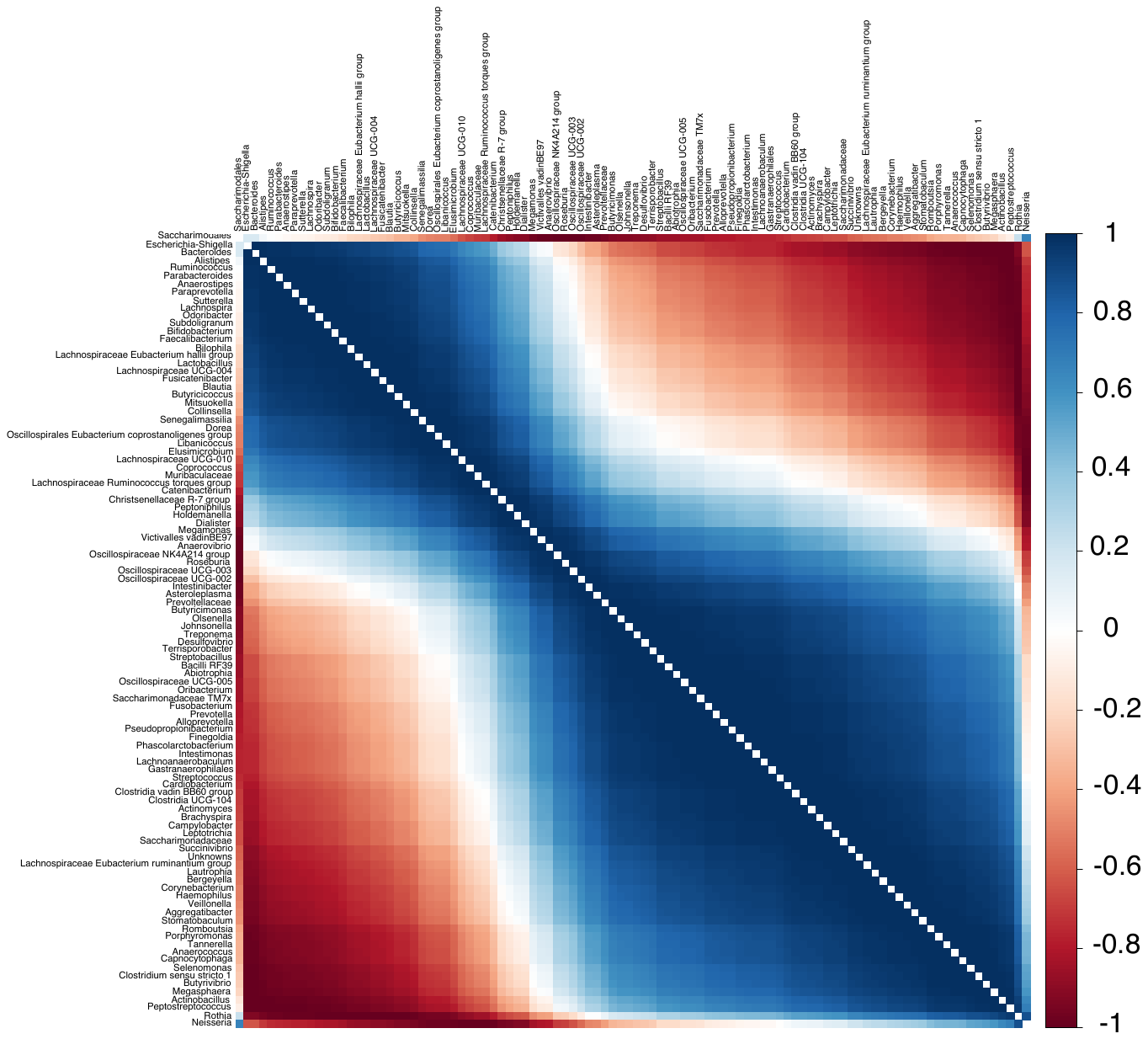

**Supplementary Figure 3**. Co-occurrence relationship between microbes recovered from the residual covariance matrix $\boldsymbol{\Sigma}$ that are not explained by the observed covariates in the GLLVM model in Figure 4. Here, blue represent the positive correlation (taxa 1 increasing in abundance leads to taxa 2 increasing in abundance), and red represent the negative relationship (taxa 1 increasing in abundance leads to taxa 2 decreasing in abundance).

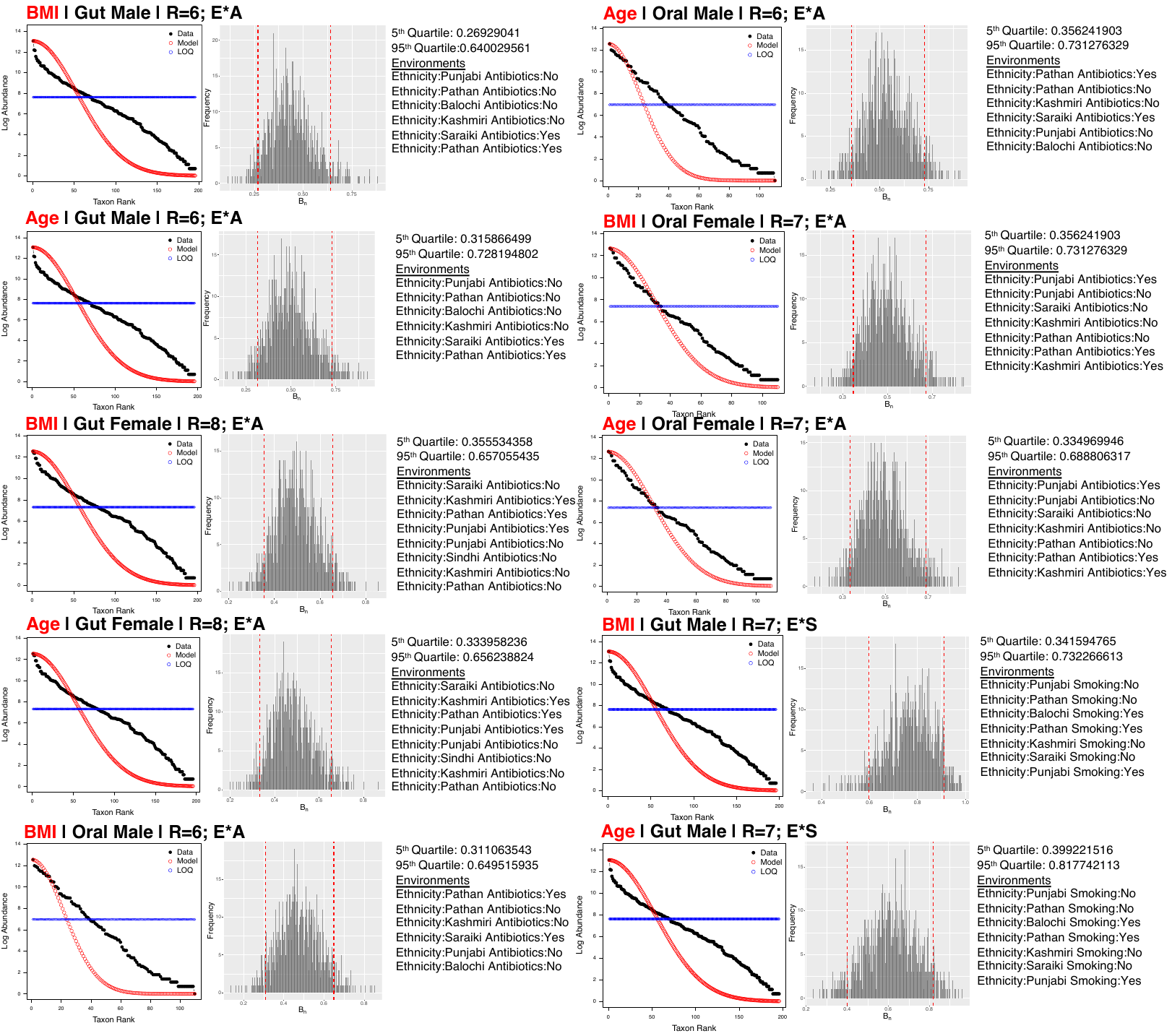

**Supplementary Figure 4**. We have applied Hurlbert's B_N_ to two environmental properties, *Age*, and *BMI* considered in this study depending on the expansive set of environments the microbes are observed in, e.g., “*BMI | Gut Male | R = 6; E * A”* represent *BMI* as the environmental property and samples from the male gut with 6 environments as a combination of *Ethnicity* (E) and *Antibiotic status* (A). Similarly, *E * S* represent all possible combinations of *Ethnicity* and *Smoking* status. All the analyses are given as three figure tuples. The left figures represent the rank distribution of the taxa observed in the dataset represented as black. The lognormal rank distribution model is then shown as red circles. The limit of quantification threshold, are then shown as blue circles, is 1.65 standard deviations from zero. Any taxa that fall below the limit of quantification were excluded from the analyses. The middle figure represents the null model distributions generated from applying Hurlbert's B_N_ calculated from 999 randomly generated taxon distributions. Red dotted lines indicate the fifth and 95th quantiles. Taxa that are high when an environmental property (*Age* or *BMI*) is low have a Hurlbert's B_N_ below the 5th quantile, and conversely, taxa that are high when the environmental property is high have a Hurlbert's B_N_ above the 95th quantile of those null models. The right figure represents the values of fifth and 95th quantiles along with the details of all possible environments.

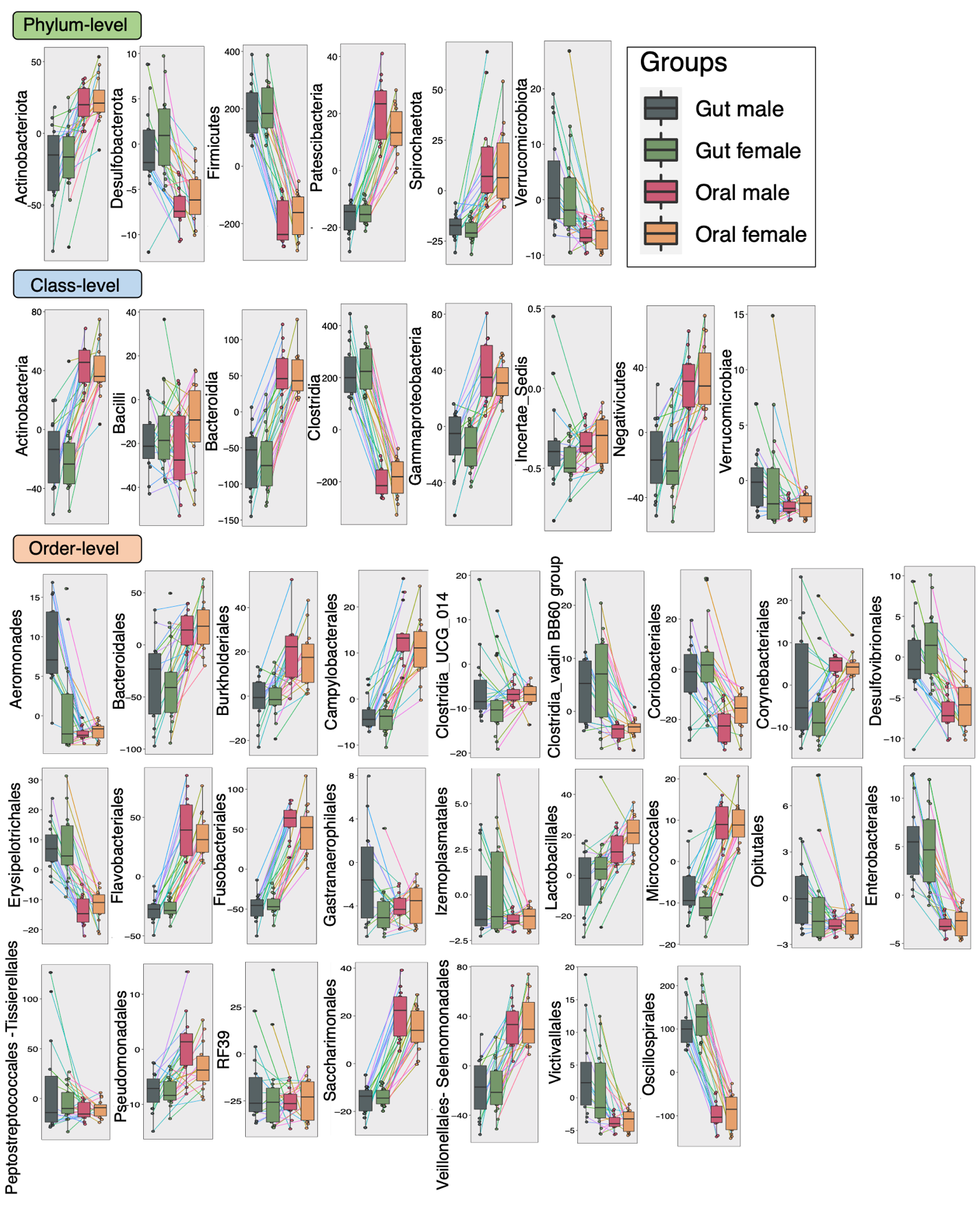

**Supplementary Figure 5**. Subset of taxa (at phylum, class, and order level) that are differentially abundant between the cohorts considered in this study using QCAT-C association test that takes into account paired nature of samples i.e., originating from the same subject connected by lines. The values represent the TSS+CLR normalized abundances of individual taxa.

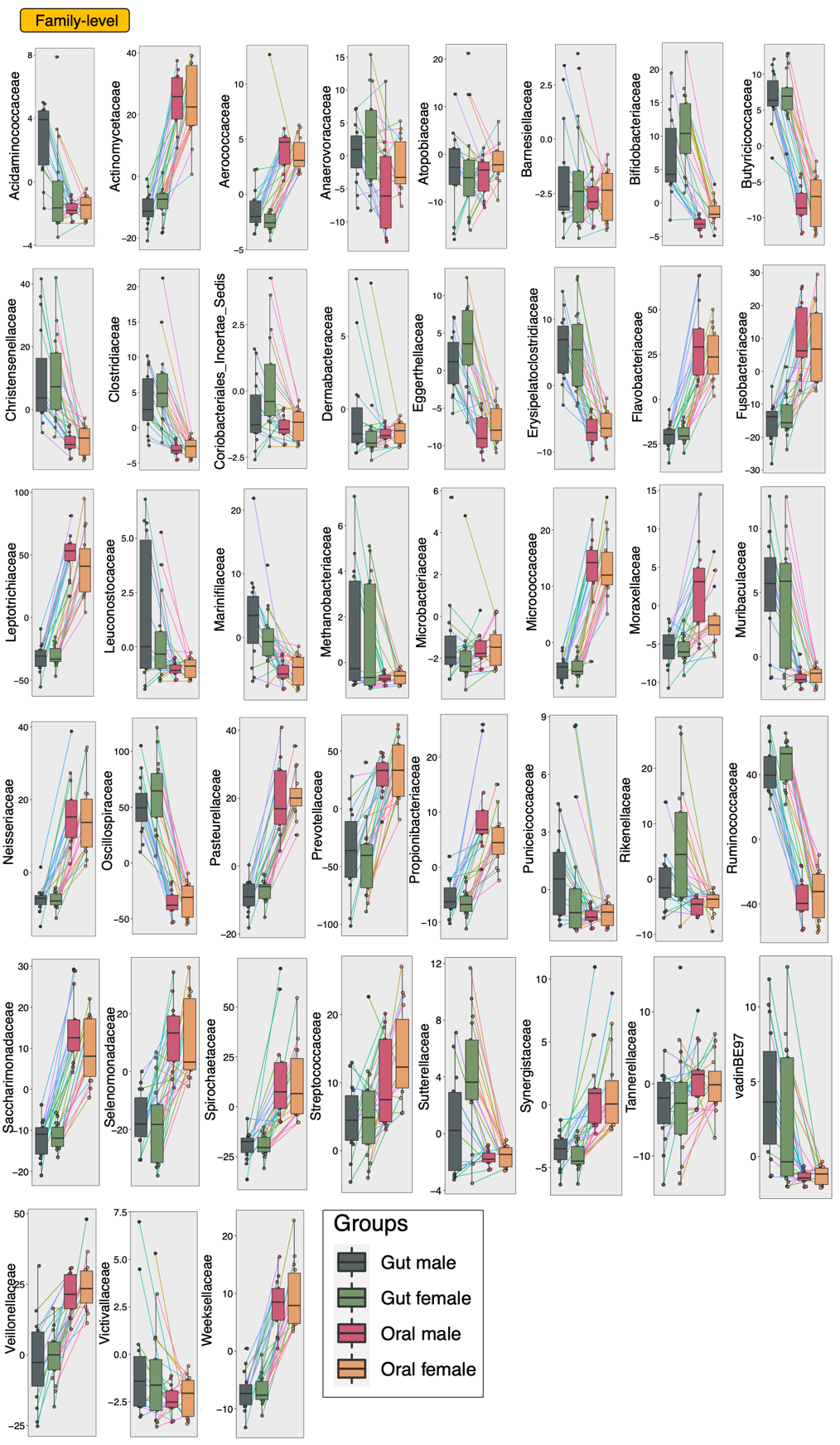

**Supplementary Figure 6**. Subset of taxa (at family level) that are differentially abundant between the cohorts considered in this study using QCAT-C association test that takes into account paired nature of samples i.e., originating from the same subject connected by lines. The values represent the TSS+CLR normalized abundances of individual taxa.

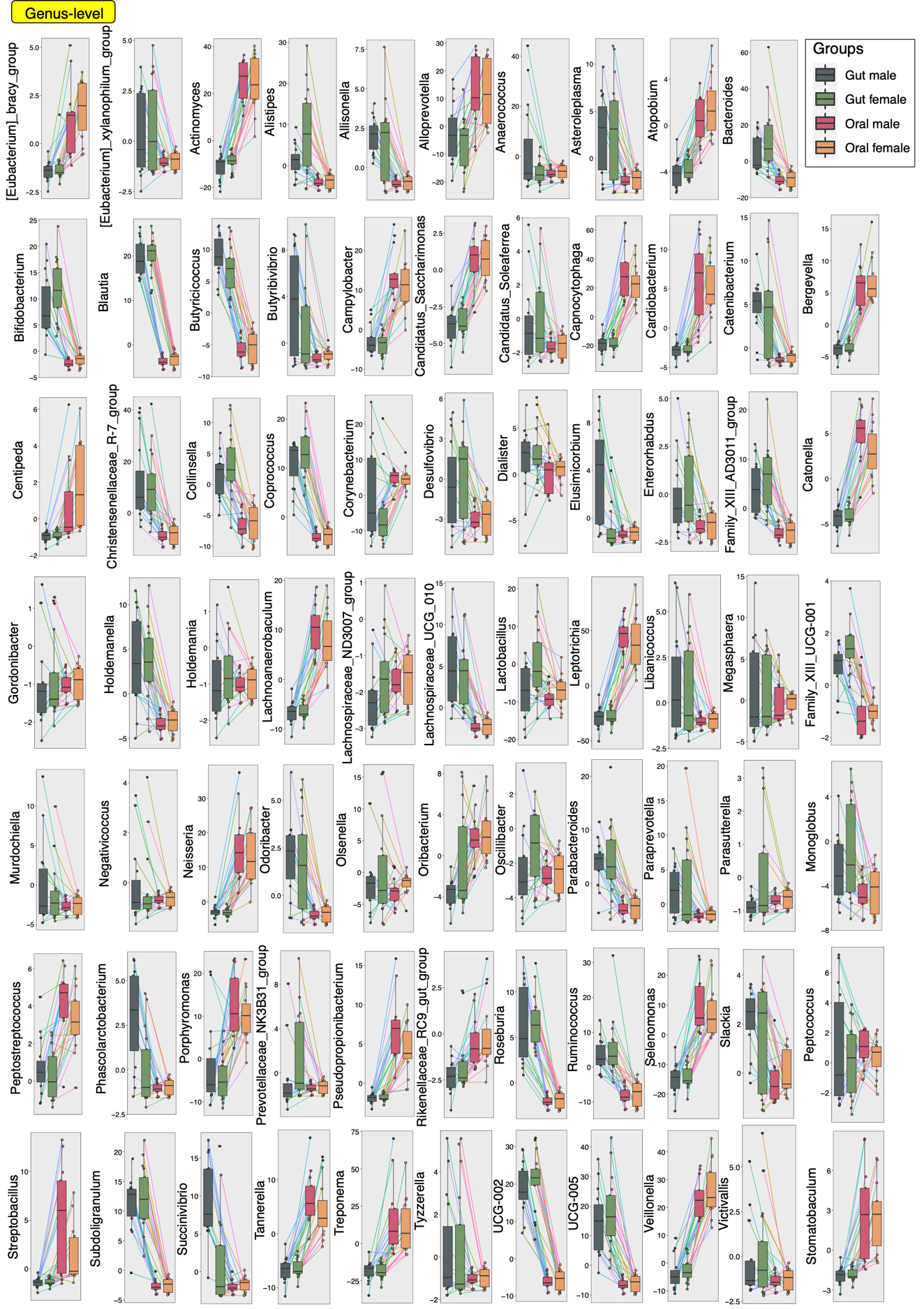

**Supplementary Figure 7**. Subset of taxa (at genus level) that are differentially abundant between the cohorts considered in this study using QCAT-C association test that takes into account paired nature of samples i.e., originating from the same subject connected by lines. The values represent the TSS+CLR normalized abundances of individual taxa.

**Supplementary Table 1.** Master list of AMR KEGG Orthologs (KOs) from <https://www.genome.jp/kegg/annotation/br01600.html>. Those AMR KOs that were detected in our study are highlighted in grey along with annotation information shown.

| **Sr No:** | **Class** | **KO** | **Name** | **Threat_level** | **Drug_group** |
| --- | --- | --- | --- | --- | --- |
| 1 | A 2b | K18698 | beta-lactamase class A TEM | A3 B1 B4 B6 B7 B8 B12 | Extended-spectrum cephalosporin (DG01776, DG01777),Monobactam (DG01454) |
| 2 | A 2b | K18699 | beta-lactamase class A SHV | B1 B4 B5 | Extended-spectrum cephalosporin (DG01776, DG01777),Monobactam (DG01454) |
| 3 | A 2b | K18767 | beta-lactamase class A CTX-M | B1 B4 B7 B8 B9 | Extended-spectrum cephalosporin (DG01776, DG01777),Monobactam (DG01454) |
| 4 | A 2b | K18797 | beta-lactamase class A PER | B1 B4 B6 B7 B8 | Extended-spectrum cephalosporin (DG01776, DG01777),Monobactam (DG01454) |
| 5 | A 2b | K19097 | beta-lactamase class A VEB | B1 B4 B6 | Extended-spectrum cephalosporin (DG01776, DG01777) |
| 6 | A 2b | K19317 | beta-lactamase class A BEL | B6 | Extended-spectrum cephalosporin (DG01776, DG01777) |
| 7 | A 2b | K18796 | beta-lactamase class A LAP |  |  |
| 8 | A 2f | K18768 | beta-lactamase class A KPC | A2 B1 B6 | Carbapenem (DG01458) |
| 9 | A 2f | K18970 | beta-lactamase class A GES | B1 B4 B6 | Extended-spectrum cephalosporin (DG01776, DG01777),Carbapenem (DG01458) |
| 10 | A 2f | K19316 | beta-lactamase class A IMI/SME | A2 | Carbapenem (DG01458) |
| 11 | A 2f | K22346 | beta-lactamase class A SME | A2 | Carbapenem (DG01458) Second-generation cephalosporin (DG01775) Monobactam (DG01454) |
| 12 | A 2c | K18795 | beta-lactamase class A CARB-1 | B6 B7 | Carbenicillin (DG00519) |
| 13 | A 2c | K19218 | beta-lactamase class A CARB-5 | B1 | Carbenicillin (DG00519) |
| 14 | A 2c | K19217 | beta-lactamase class A CARB-17 |  |  |
| 15 | D 2d | K18794 | beta-lactamase class D OXA-51 | B1 | Carbapenem (DG01458) |
| 16 | D 2d | K19318 | beta-lactamase class D OXA-213 | B1 | Carbapenem (DG01458) |
| 17 | D 2d | K18971 | beta-lactamase class D OXA-24 | B1 | Carbapenem (DG01458) |
| 18 | D 2d | K18793 | beta-lactamase class D OXA-23 | A2 B1 | Carbapenem (DG01458) |
| 19 | D 2d | K19319 | beta-lactamase class D OXA-134 | B1 | Carbapenem (DG01458) |
| 20 | D 2d | K19320 | beta-lactamase class D OXA-211 | B1 | Carbapenem (DG01458) |
| 21 | D 2d | K19321 | beta-lactamase class D OXA-214 | B1 | Extended spectrum penicillin (DG01780) Carbapenem (DG01458) (weak) |
| 22 | D 2d | K19322 | beta-lactamase class D OXA-229 | B1 | Carbapenem (DG01458) |
| 23 | D 2d | K18972 | beta-lactamase class D OXA-58 | B1 | Carbapenem (DG01458) |
| 24 | D 2d | K21266 | beta-lactamase class D OXA-286 | B1 |  |
| 25 | D 2d | K18973 | beta-lactamase class D OXA-50 | B6 | Narrow-spectrum penicillin (DG01779) |
| 26 | D 2d | K19211 | beta-lactamase class D OXA-62 |  | Carbapenem (DG01458) |
| 27 | D 2d | K18791 | beta-lactamase class D OXA-2 | B1 B4 B6 B7 B8 | Extended-spectrum cephalosporin (DG01776, DG01777) |
| 28 | D 2d | K18792 | beta-lactamase class D OXA-10 | B1 B4 B6 | Extended-spectrum cephalosporin (DG01776, DG01777) |
| 29 | D 2d | K18976 | beta-lactamase class D OXA-48 | A2 | Carbapenem (DG01458 |
| 30 | D 2d | K19210 | beta-lactamase class D OXA-61 | B2 | Narrow-spectrum penicillin (DG01779) |
| 31 | D 2d | K19212 | beta-lactamase class D OXA-63 |  | Narrow-spectrum penicillin (DG01779) |
| 32 | D 2d | K18790 | beta-lactamase class D OXA-1 | B4 B6 B7 B9 | Extended-spectrum cephalosporin (DG01776, DG01777),Extended spectrum penicillin (DG01780) |
| 33 | D 2d | K19098 | beta-lactamase class D OXA-9 |  | Narrow-spectrum penicillin (DG01779) |
| 34 | D 2d | K19209 | beta-lactamase class D OXA-42 |  | Narrow-spectrum penicillin (DG01779) |
| 35 | D 2d | K19213 | beta-lactamase class D OXA-12 |  | Narrow-spectrum penicillin (DG01779) |
| 36 | D 2d | K21276 | beta-lactamase class D OXA-22 |  | Narrow-spectrum penicillin (DG01779) |
| 37 | D 2d | K21277 | beta-lactamase class D OXA-60 |  | Narrow-spectrum penicillin (DG01779),Carbapenem (DG01458) (weak) |
| 38 | D 2d | K22331 | beta-lactamase class D OXA-184 | B2 |  |
| 39 | D 2d | K22332 | beta-lactamase class D OXA-548 |  |  |
| 40 | D 2d | K22333 | beta-lactamase class D OXA-493 |  |  |
| 41 | D 2d | K22334 | beta-lactamase class D OXA-464 |  |  |
| 42 | D 2d | K22335 | beta-lactamase class D OXA-114 |  | Extended spectrum penicillin (DG01780),Third-generation cephalosporin (DG01776) |
| 43 | D 2d | K22351 | beta-lactamase class D OXA-209 |  | Extended-spectrum penicillin (DG01780) |
| 44 | D 2d | K22352 | beta-lactamase class D OXA-29 |  | Extended-spectrum penicillin (DG01780) |
| 45 | C 1 | K19095 | beta-lactamase class C CMY-1 | B4 | Extended-spectrum cephalosporin (DG01776, DG01777) |
| 46 | C 1 | K19096 | beta-lactamase class C CMY-2 | B4 B7 | Second-generation cephalosporin (DG01775),Third-generation cephalosporin (DG01776) |
| 47 | C 1 | K19100 | beta-lactamase class C DHA |  |  |
| 48 | C 1 | K19101 | beta-lactamase class C FOX |  |  |
| 49 | C 1 | K19214 | beta-lactamase class C ACC |  |  |
| 50 | C 1 | K19215 | beta-lactamase class C ACT/MIR |  | Extended-spectrum penicillin (DG01780),Second-generation cephalosporin (DG01775) |
| 51 | C 1 | K20319 | beta-lactamase class C ADC |  |  |
| 52 | C 1 | K20320 | beta-lactamase class C PDC |  |  |
| 53 | B | K18782 | metallo-beta-lactamase class B IMP | A2 B1 B4 B6 B9 | Extended-spectrum cephalosporin (DG01776, DG01777),Carbapenem (DG01458) |
| 54 | B | K18781 | metallo-beta-lactamase class B VIM | B4 B6 | Extended-spectrum cephalosporin (DG01776, DG01777),Carbapenem (DG01458) |
| 55 | B | K18780 | metallo-beta-lactamase class B NDM | A2 B1 | Carbapenem (DG01458) |
| 56 | B | K19099 | metallo-beta-lactamase class B GIM | A2 B6 | Carbapenem (DG01458) |
| 57 | B | K19216 | metallo-beta-lactamase class B IND |  | Carbapenem (DG01458) |
| 58 | O | K17840 | aminoglycoside 2'-N-acetyltransferase I | B6 B12 | Aminoglycoside (DG01447) |
| 59 | O | K03395 | aminoglycoside 3-N-acetyltransferase I | B1 B6 B7 | Aminoglycoside (DG01447) |
| 60 | O | K19275 | aminoglycoside 3-N-acetyltransferase II | B1 B7 | Aminoglycoside (DG01447) |
| 61 | O | K19276 | aminoglycoside 3-N-acetyltransferase IV | B6 | Aminoglycoside (DG01447) |
| 62 | O | K19277 | aminoglycoside 3-N-acetyltransferase VI | B7 | Aminoglycoside (DG01447) |
| 63 | O | K19278 | aminoglycoside 6'-N-acetyltransferase Ib | B1 B6 | Aminoglycoside (DG01447) |
| 64 | O | K19301 | aminoglycoside 6'-N-acetyltransferase II | B1 B6 | Aminoglycoside (DG01447) |
| 65 | O | K18815 | aminoglycoside 6'-N-acetyltransferase I | B1 B6 B7 | Aminoglycoside (DG01447) |
| 66 | O | K18816 | aminoglycoside 6'-N-acetyltransferase I | B1 B6 B7 B11 | Aminoglycoside (DG01447) |
| 67 | O | K17881 | aminoglycoside 2''-adenylyltransferase | B1 | Aminoglycoside (DG01447) |
| 68 | O | K19544 | aminoglycoside 4'-adenylyltransferase | B6 | Aminoglycoside (DG01447) |
| 69 | O | K19272 | aminoglycoside 3'-phosphotransferase I | B1 B7 | Aminoglycoside (DG01447) |
| 70 | O | K19300 | aminoglycoside 3'-phosphotransferase II | B6 | Aminoglycoside (DG01447) |
| 71 | O | K19299 | aminoglycoside 3'-phosphotransferase III | B11 | Aminoglycoside (DG01447) |
| 72 | O | K19274 | aminoglycoside 3'-phosphotransferase VI | B1 | Aminoglycoside (DG01447) |
| 73 | O | K19315 | aminoglycoside 3'-phosphotransferase XV | B6 | Aminoglycoside (DG01447) |
| 74 | O | K10673 | streptomycin 3"-kinase | B1 B6 B7 | Aminoglycoside (DG01447) |
| 75 | O | K04343 | streptomycin 6-kinase | B1 B6 B7 | Aminoglycoside (DG01447) |
| 76 | O | K18845 | 16S rRNA (guanine(1405)-N(7))-methyltransferase | B1 B6 | Aminoglycoside (DG01447) |
| 77 | O | K18220 | ribosomal protection tetracycline resistance protein | A1 | Tetracycline (DG00005) |
| 78 | O | K00561 | 23S rRNA (adenine-N6)-dimethyl transferase | B11 | Macrolide antibiotic (DG01551) |
| 79 | O | K18231 | macrolide transport system ATP-binding/permease protein | B1 B11 | Macrolide antibiotic (DG01551) |
| 80 | O | K06979 | macrolide phosphotransferase | B1 | Macrolide antibiotic (DG01551) |
| 81 | O | K08217 | MFS transporter, DHA3 family, macrolide efflux protein | B7 B11 | Macrolide antibiotic (DG01551) |
| 82 | O | K18552 | MFS transporter, DHA1 family, florfenicol/chloramphenicol resistance protein | B6 B7 | Phenicol (DG01576) |
| 83 | O | K19271 | chloramphenicol O-acetyltransferase type A | A3 B1 B6 B7 B11 | Phenicol (DG01576) |
| 84 | O | K00638 | chloramphenicol O-acetyltransferase type B | B1 B6 B7 | Phenicol (DG01576) |
| 85 | O | K18554 | chloramphenicol 3-O phosphotransferase | B12 | Phenicol (DG01576) |
| 86 | O | K18589 | dihydrofolate reductase DfrA | B1 B6 B7 | Trimethoprim (DG01581) |
| 87 | O | K19643 | dihydrofolate reductase DfrA | B1 | Trimethoprim (DG01581) |
| 88 | O | K18590 | dihydrofolate reductase DfrA | B7 | Trimethoprim (DG01581) |
| 89 | O | K19644 | dihydrofolate reductase DfrA | B7 | Trimethoprim (DG01581) |
| 90 | O | K19645 | dihydrofolate reductase DfrB | B6 B7 | Trimethoprim (DG01581) |
